# SOX9 acts as a dynamic pioneer factor inducing stable changes in the chromatin landscape to reprogram endothelial cells

**DOI:** 10.1101/2020.09.11.293993

**Authors:** Bettina M. Fuglerud, Sibyl Drissler, Jeremy Lotto, Tabea L. Stephan, Avinash Thakur, Rebecca Cullum, Pamela A. Hoodless

## Abstract

The transcription factor SOX9 is expressed in multiple tissues during embryogenesis and directs developmental processes. SOX9 is activated upon endothelial-to-mesenchymal transition (EndMT) in the developing heart, but its role in regulating this process is less clear. Using human umbilical vein endothelial cells as an EndMT model, we show that SOX9 expression alone is sufficient to activate mesenchymal enhancers and steer endothelial cells towards a mesenchymal fate. By genome-wide mapping of the chromatin landscape, we show that SOX9 acts as a pioneer transcription factor, having the ability to open chromatin and lead to deposition of active histone marks at a specific subset of previously silent enhancers, guided by SOX motifs and H2A.Z enrichment. This leads to a switch in enhancer activity states resulting in activation of mesenchymal genes and concurrent suppression of endothelial genes to drive EndMT. Moreover, we show that SOX9 chromatin binding is dynamic, but induces stable changes in the chromatin landscape. Our data also show widespread SOX9 chromatin scanning in silent chromatin that is not associated with SOX motifs or H2A.Z enrichment. Our study highlights the crucial developmental role of SOX9 and provides new insight into key molecular functions of SOX9 in the chromatin landscape and mechanisms of EndMT.

**Graphical abstract:** 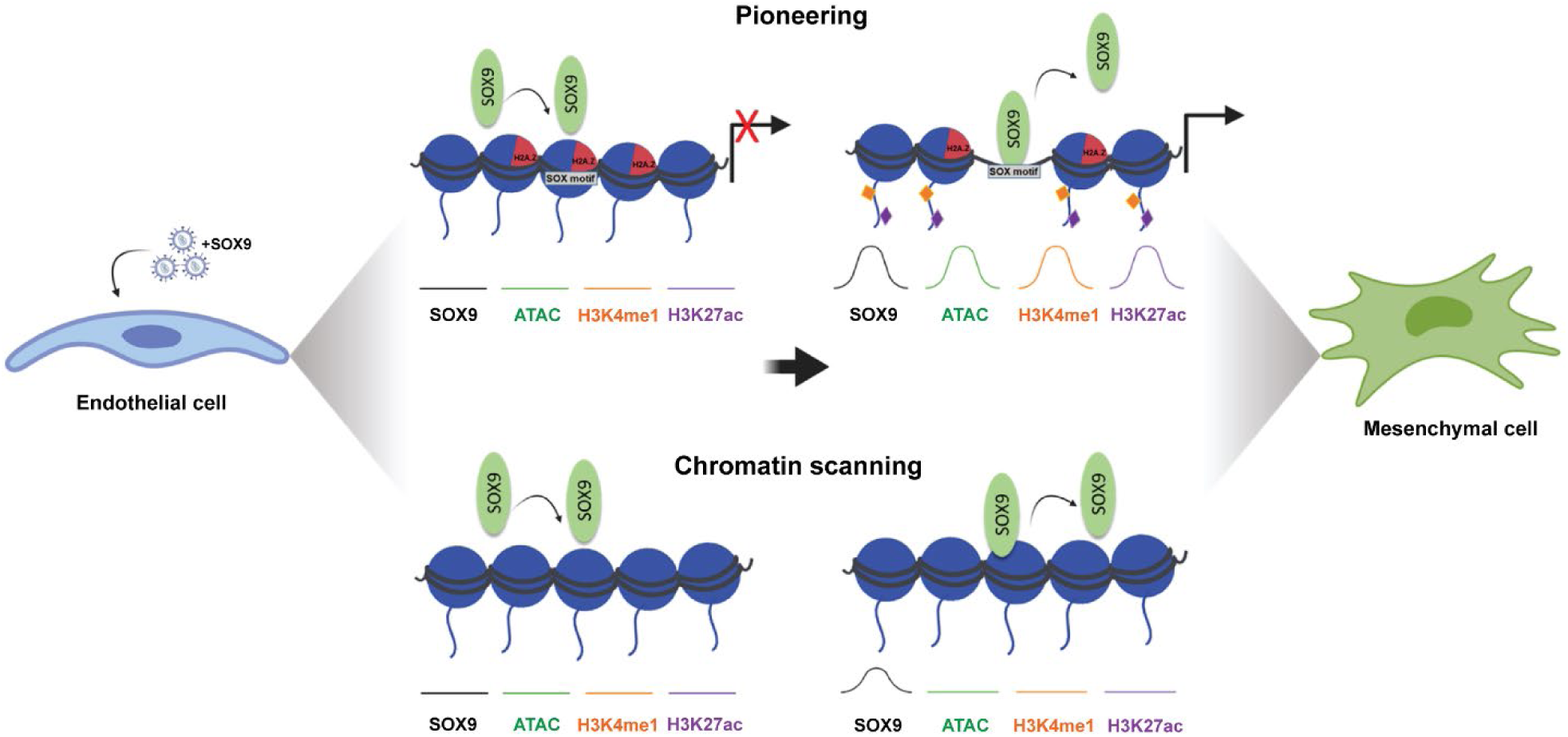

## Introduction

During embryogenesis, SOX9 directs developmental processes including chondrogenesis, gliogenesis, sex determination, as well as development of the heart valves, hair follicles, pancreas, and liver (summarized in (Jo et al. 2014)). SOX9 is a member of the SOX family of transcription factors (TFs), which contains a highly conserved high mobility group (HMG) domain binding to ACAAa/tG-like DNA sequences. The HMG domain was first identified in *SRY*, a crucial factor for mammalian sex determination (Sinclair et al. 1990; Harley, Lovell-Badge, and Goodfellow 1994). Around 20 SOX proteins with divergent developmental functions have been identified, of which many act as master regulators that control critical transcriptional programs to determine cell fate and differentiation. SOX9, and other SOX factors, bind DNA as either a monomer or a homo- or heterodimer and this can alter functionality (Hou, Srivastava, and Jauch 2017).

Most of the understanding regarding biological functions of SOX9 has come from studies involving chondrogenesis, which have revealed its role in regulating extracellular matrix (ECM) production and organization (Bi et al. 1999; Hanley et al. 2008). During normal development, SOX9 is also crucial for processes involving epithelial-to-mesenchymal transition (EMT) and the closely related process endothelial-to-mesenchymal transition (EndMT) (Lincoln et al. 2007; Akiyama et al. 2004; M. Cheung et al. 2005). Given the significance of EMT as a disease mechanism in cancer progression, SOX9 can drive cancer invasion and metastasis in multiple cancers, including prostate, breast, colon, gastric, and lung (summarized in (Aguilar-Medina et al. 2019)). SOX9 is also involved in promoting liver and cardiac fibrosis by stimulating inappropriate ECM deposition (Lee and Friedman 2011; Lacraz et al. 2017). SOX9 is essential during cushion formation in cardiac valve development, which involves EndMT, and where SOX9 activates transcription of key regulators of heart valve development (Akiyama et al. 2004; Garside et al. 2015). However, the role of SOX9 in EndMT induction has not been evident.

At the molecular level, SOX9 has been suggested to alter epigenetic signatures, but most of the work has been done on a single locus in the *COL2A1* enhancer. SOX9 remodeled chromatin and activated transcription at this site using an *in vitro* assembled chromatin template (Coustry et al. 2010). At this same enhancer, the histone acetyltransferase, p300, and SOX9 synergistically acetylated histones on a chromatin template, while SOX9 was not required to assist p300 in acetylating free histones (Furumatsu et al. 2005). The latter strongly suggests that SOX9 recruits the histone modifier to the chromatinized template. In chondrogenic systems, SOX9 can activate enhancers (Cheung et al. 2020), but may not be required for the initiation of chromatin changes (Liu et al. 2018).

Some developmental TFs have the remarkable ability to reprogram one cell type into another by engaging genes that are developmentally silenced and in closed chromatin, thus acting as pioneer TFs to initiate transcriptional events (Iwafuchi-Doi and Zaret 2016). In hair follicle stem cells, SOX9 has been put forward as a pioneer TF (Adam et al. 2015); however, this study only showed increased activation of enhancers and did not identify whether SOX9 opens chromatin, a hallmark of pioneer TFs. SOX9 acts as a master regulator in heart valve development, binding regulatory regions that control TFs important for proper valve development, but its impact on the chromatin landscape is not yet characterized (Garside et al. 2015). In summary, the epigenetic requirements of SOX9 to alter chromatin and drive cell fate decisions remains uncertain and may vary between cell types.

In the present work, we have used human umbilical vein endothelial cells (HUVECs) to study cell reprogramming initiated by SOX9 and its role in restructuring the chromatin landscape.

Transcriptome sequencing revealed that ectopic expression of SOX9 in endothelial cells is sufficient to induce expression of mesenchymal marker genes. In addition, the cells acquired increased invasive properties and a mesenchymal morphology. We used the chromatin profiling methods CUT&RUN and CUT&Tag to map SOX9 binding and histone marks (H3K4me1, H3K27ac, and H3K27me3). In addition, we mapped chromatin accessibility by ATAC-seq.

Altogether, we found that at a subset of regions that are silent in unstimulated HUVEC, SOX9 occupancy increases chromatin accessibility and enrichment of active histone marks to induce gene expression, demonstrating SOX9’s ability to act as a pioneer TF to reprogram cell fate. This pioneer function is motif encoded and occurs predominantly in distal regulatory regions with enrichment of the histone variant H2A.Z. Furthermore, we show that at most pioneering sites, SOX9 binding is highly dynamic, but nevertheless causes stable changes in the chromatin landscape. Together, our work demonstrates how a single TF can have widespread effects on chromatin structure and reprogram cell fate.

## Results

### SOX9 expression in endothelial cells promotes a mesenchymal phenotype

Mouse embryonic heart valve formation initiates when endothelial cells of the heart, endocardial cells, transition into mesenchymal cells and migrate to populate the cardiac cushions, the precursors of the cardiac valves. To determine when SOX9 expression is initiated during EndMT, we examined SOX9 levels in mouse cardiac cushions. At embryonic day (E) 9.5, the endothelial master TF ERG is expressed in all endocardial cells and downregulated within emerging mesenchymal cells, which express SOX9 as they undergo EndMT and populate the cardiac cushions (Fig.1A). Interestingly, many endocardial cells lining the cushions express both master regulators. This suggests that SOX9 expression is induced in endothelial cells lining the cardiac cushions, prior to overt EndMT, and continues to be expressed through the transition into migrating mesenchymal cells. By E10.5, we detected fewer cells co-expressing SOX9 and ERG, and the SOX9-expressing mesenchymal cells had proliferated, populating the cardiac cushion. By E12.5, we observed a further expansion of the mesenchyme and detected high levels of the ECM protein Periostin (POSTN) and the intermediate filament protein Vimentin (VIM), characteristic of emergent mesenchymal identity during endocardial EndMT.

**Figure 1.**
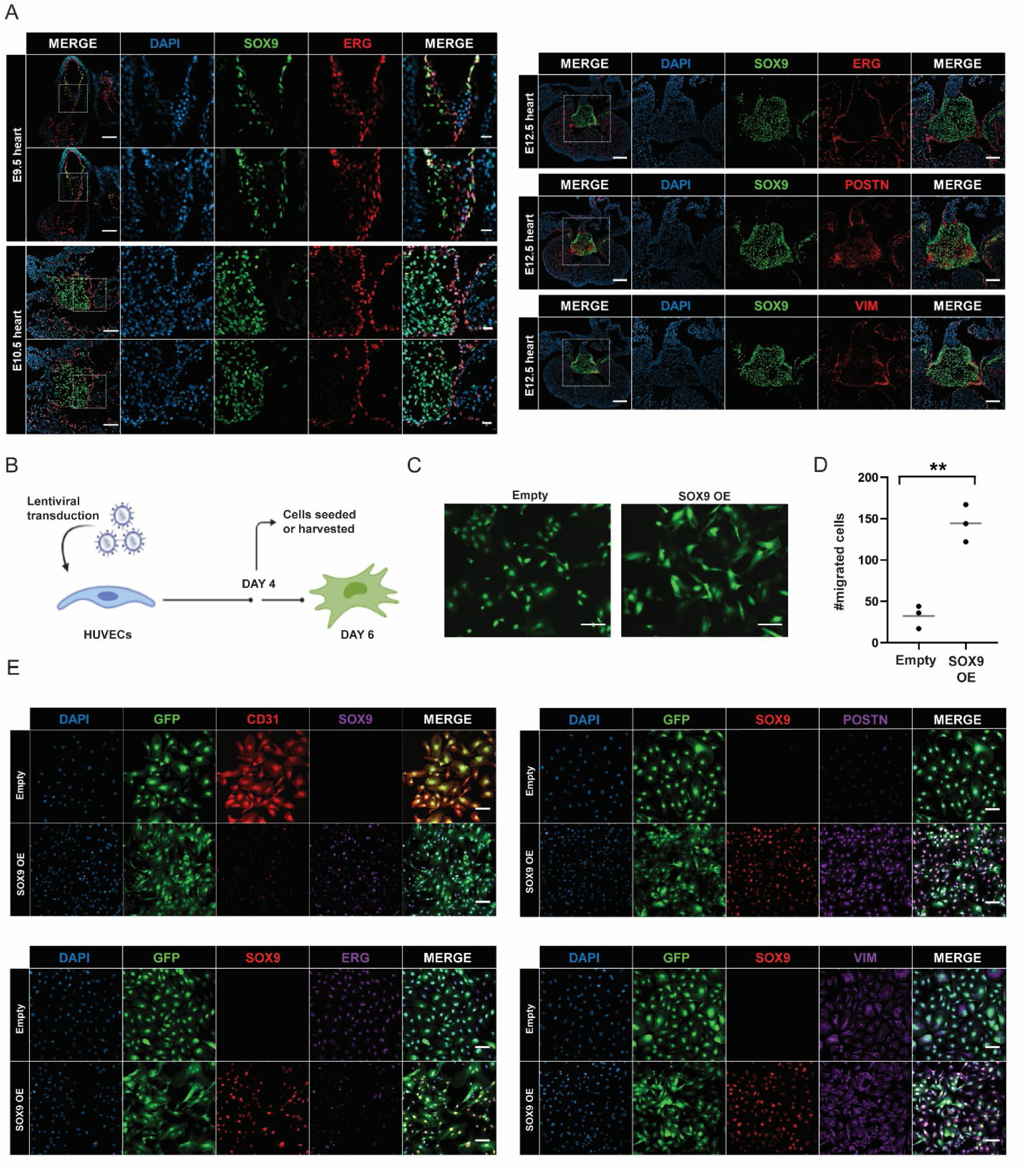
SOX9 is expressed at the onset of EndMT in endothelial cells in mouse embryonic hearts and promotes a mesenchymal phenotype of human endothelial cells. A) SOX9, ERG, Periostin (POSTN), and Vimentin (VIM) immunostaining of mouse embryonic hearts at embryonic day (E) 9.5,10.5 and 12.5. Scale bar whole hearts, 100 µm. Scale bar areas of interest, 25 µm. B) Schematic overview of experimental setup. The figure was created with Biorender. C) Fluorescence microscopy of the HUVECs six days after transduction with SOX9 or empty vector plasmid, both containing a GFP gene, demonstrating the change in morphology with SOX9 expression. Scale bars, 100 μm. D) Indicated number of HUVECs migrated after transduction with SOX9 or empty vector. Three transwells were used per condition. Significance was evaluated by unpaired, two-tailed t-tests with P-values (*P<0.05; **P<0.01; ***P<0.001; ****P<0.0001; ns P>0.05). Mean is indicated by gray line. E) SOX9, ERG, CD31, Periostin, and Vimentin (VIM) immunostaining of HUVECs six days after transduction with SOX9 or empty vector. Scale bars, 100 µm.

To overcome the challenge of isolating SOX9-expressing cells from E9.5 cushions and to determine if ectopic SOX9 expression in endothelial cells would force cells to adopt a mesenchymal phenotype using a human model, we used HUVECs, a recognized EndMT model that lacks endogenous SOX9 expression. we transduced HUVECs with constructs encoding human SOX9 or an empty vector control, both containing a green fluorescent protein (GFP) reporter gene (Fig. 1B). Expression of SOX9 was confirmed (Fig. S1A) and cellular GFP levels were monitored. By six days post-transduction the cellular morphology had changed from the cobblestone-like shape of endothelial cells to an elongated spindle-shape characteristic of mesenchymal cells (Fig. 1C). This change in morphology is consistent with other studies’ observations of EndMT of HUVECs (Yu et al. 2017; Liguori et al. 2019; Noseda et al. 2004).

Notably, we could continue to passage the cells without the cells transitioning back to their original morphology, suggesting the mesenchymal morphology induced by SOX9 was permanent and robust. As expected from mesenchymal cells, the SOX9-expressing cells displayed increased migratory properties (Fig. 1D). To further characterize the SOX9-expressing cells, we immunostained cells for proteins characteristic for endothelial and mesenchymal cells (Fig. 1E). We observed a clear downregulation of the endothelial surface protein CD31 as well as ERG in SOX9-expressing cells compared to cells without SOX9. SOX9-expressing cells also had high Periostin levels, while Vimentin levels were similar in both cell types. Of note, while Vimentin is present in most cell types and higher in mesenchymal cells, Periostin is more specific to mesenchymal cells (Thul et al. 2017).

The above results suggest that SOX9 induces EndMT during embryonic heart valve development and that ectopic expression of SOX9 alone in endothelial cells is sufficient for direct reprogramming to a mesenchymal cell fate.

### SOX9 induces a switch in enhancers and alters expression of EndMT genes

To investigate global transcriptional changes caused by ectopic SOX9 expression, we employed RNA-seq using HUVECs harvested four days after transduction with SOX9 or empty vector to assay cells in their transition towards mesenchymal cell identity. 4806 genes were differentially expressed between the two conditions (log2 ratio >0.5 or <-0.5, p-value <0.05) (Fig. S1B and S1C). Of these, 2587 genes were upregulated and 2219 genes were downregulated in cells transduced with SOX9 compared to empty vector (Fig. 2A). Among the most upregulated genes were components of ECM, including collagens, as well as mesenchymal markers like Periostin (*POSTN*), N-cadherin (*CDH2*), and Cadherin-11 (*CDH11*). The downregulated genes included endothelial markers like von Willebrand factor (*VWF*), VE-cadherin (*CDH5*), and CD31 (*PECAM1*), as well as some key endothelial-associated TFs, such as SOX7, SOX18, ERG and LMO2 (Fig. 2A, right panel). Gene Set Enrichment Analysis (GSEA) revealed that genes upregulated in response to SOX9 were significantly enriched for hallmark gene sets for EMT (Fig. 2B). Downregulated genes were enriched for inflammatory signaling pathways, reflecting the role of endothelial cells as regulators of inflammatory responses. Although EMT and EndMT are similar processes, both leading to a mesenchymal phenotype, gene sets for EndMT currently do not exist in gene ontology (GO) databases. Therefore, we generated a list of endothelial and mesenchymal genes, including genes shown to be down- or up-regulated upon EndMT (Table S1). SOX9 expression resulted in enhanced expression of mesenchymal genes, with concurrent downregulation of endothelial gene expression (Fig. 2C). Of note, we observed a downregulation of some genes encoding TFs known to have a role in EMT, like *SNAI1* and *ZEB1*, indicating that upregulation of mesenchymal markers does not depend on these TFs in this context.

**Figure 2.**
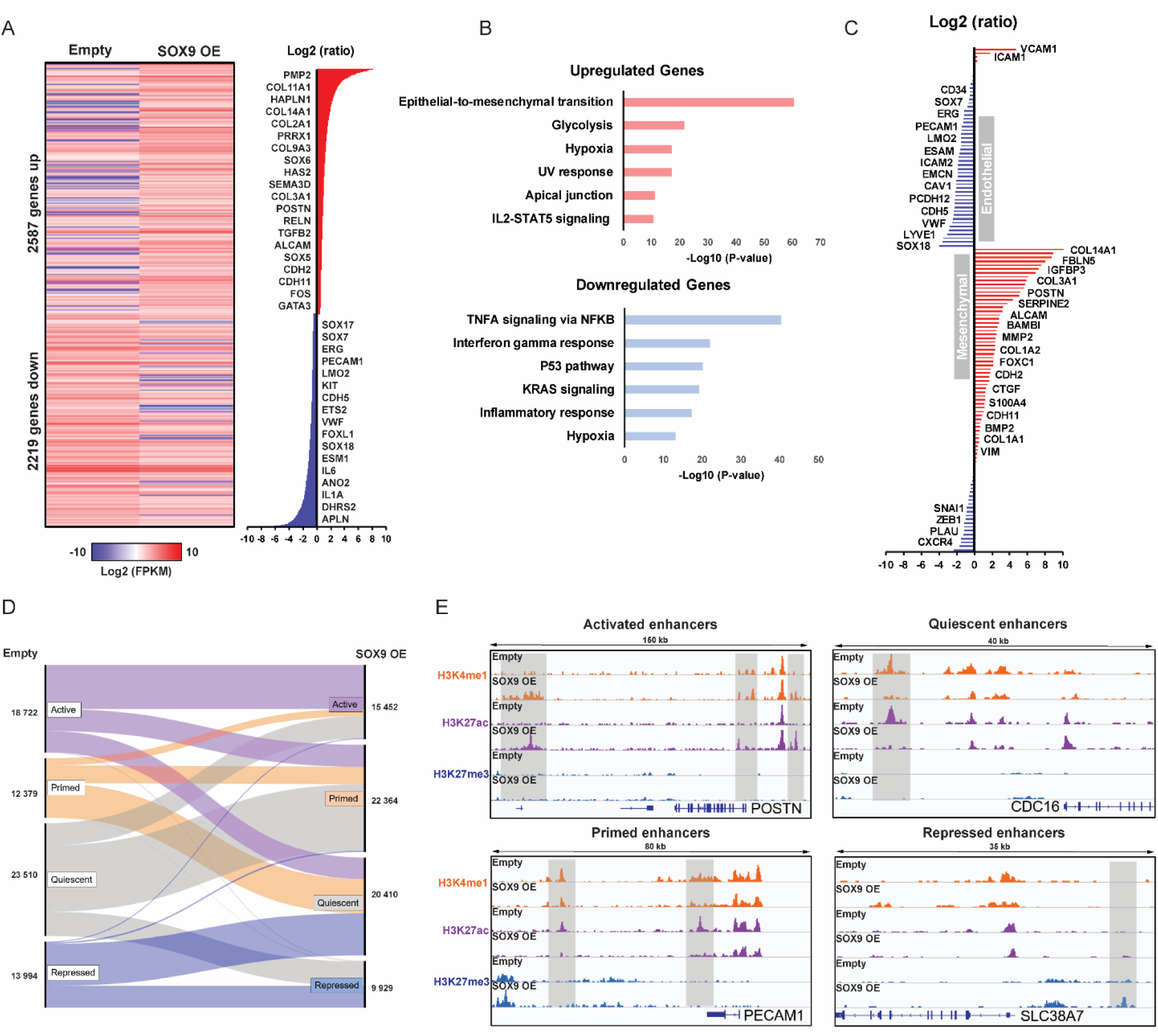
SOX9 alters expression of EndMT genes by inducing a switch in enhancers. **A)** Heatmap showing differentially expressed genes between cells transduced with SOX9 or empty vector. Heatmap shows log2 FPKM values while log2 ratios are displayed in the right panel with selected genes highlighted. Mean values of three biological replicates are shown. **B)** Top enriched Hallmark gene sets (based on P-values) for upregulated and downregulated genes upon SOX9 expression. **C)** Log2 ratios of SOX9 OE over empty vector for “endothelial” and “mesenchymal” genes with selected genes highlighted. **D)** Alluvial plot showing dynamics in enhancer states between cells transduced with SOX9 or empty vector. Alluvium colours originates from the state of the enhancers in the empty vector control cells. The number of enhancers in each alluvium is indicated on the sides. **E)** IGV browser tracks of representative loci displaying a switch in enhancer states. Dynamic enhancers are highlighted with gray boxes and the titles indicated the new state after SOX9 OE. Gene name is upstream of TSS.

Enhancers are crucial in regulating gene expression required to define cell type specificity, so we next examined the global dynamics of histone marks at enhancers after SOX9 expression. Cells were harvested four days after transduction and we used CUT&Tag to profile the histone modifications known to represent enhancers: H3K4me1, H3K27ac, and H3K27me3. We defined all enhancers as being more than 5000 bp upstream or 500 bp downstream from a transcriptional start site (TSS) and classified enhancer states as active (H3K27ac), primed (H3K4me1 only), quiescent (no marks), or repressed (H3K27me3). SOX9 expression caused a major switch in enhancer states; in total 50607 enhancers changed from one state to another (Fig. 2D and 2E). 6298 enhancers that were primed, quiescent, or repressed in endothelial cells prior to SOX9 expression, were activated with SOX9 expression. The genes associated with these activated enhancers were enriched for GO terms for mesenchymal functions, whereas the enhancers that lost active histone marks were linked with endothelial functions (Fig. S2A). In contrast, enhancers that remained active (stable enhancers) were not specifically enriched for mesenchymal processes (Fig. S2B). Together, our analysis of the transcriptome and epigenetic landscape show that SOX9 expression is sufficient to induce EndMT by causing a switch in enhancers that leads to activation of mesenchymal gene expression and downregulation of endothelial marker genes.

### CUT&RUN identifies SOX9 direct target genes

To examine the regulatory role of SOX9 in directing EndMT, we used CUT&RUN to profile SOX9 chromatin binding genome wide in HUVECs four days after transduction. We identified 3473 SOX9 bound regions not present in the background set (Fig. 3A). Approximately 6% of all SOX9 bound regions were either directly over a TSS or in the 5 kb proximal promoter regions (Fig. 3B). About 45% of bound regions were located within gene bodies, whereas 49% were distal, indicating that in HUVECs, SOX9 primarily binds distal regulatory elements inside or outside gene bodies rather than promoter regions of genes.

**Figure 3.**
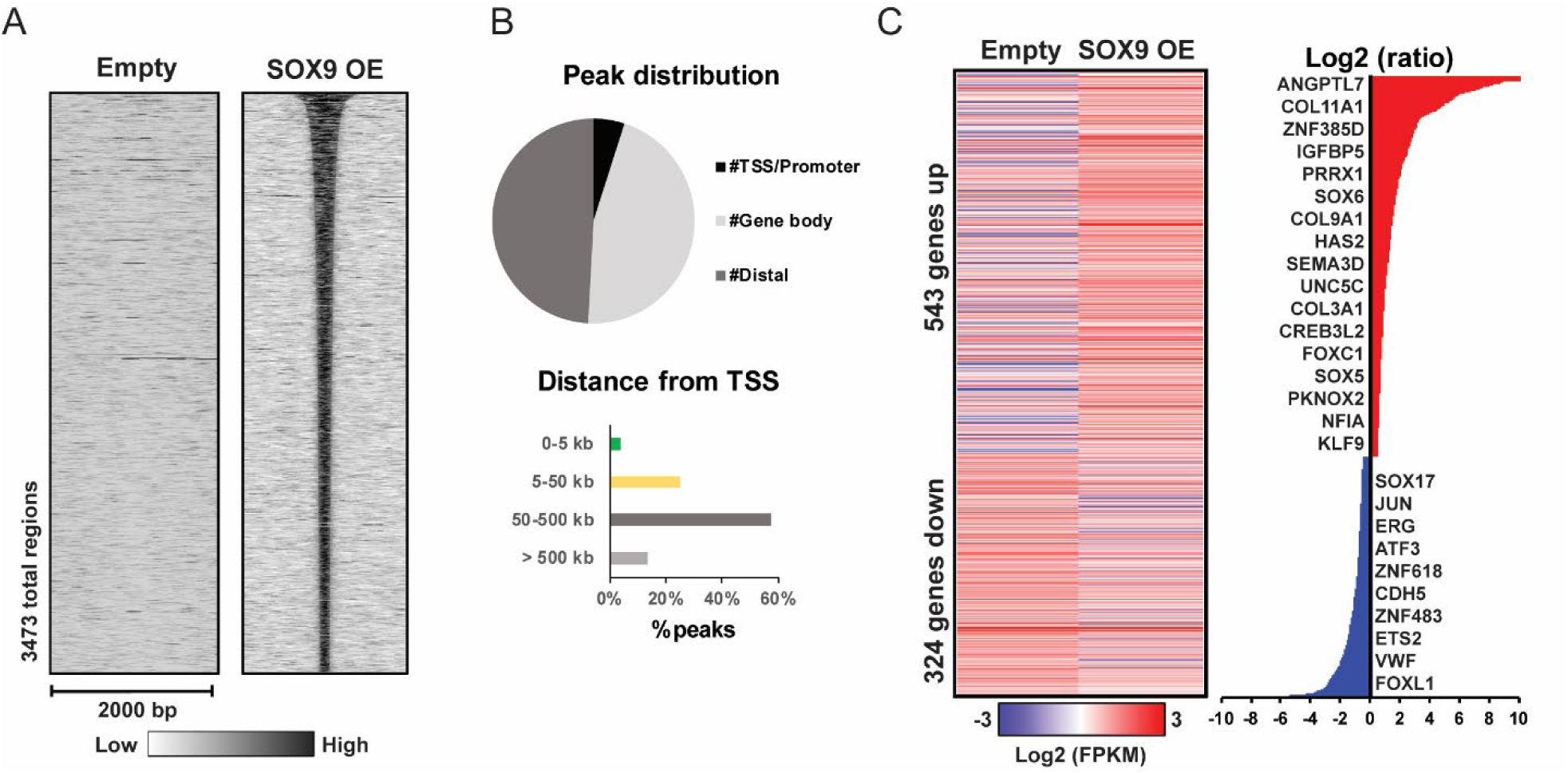
CUT&RUN profiling identifies SOX9 direct target genes. **A)** Heatmap displaying SOX9 CUT&RUN signal within a 2 kb window around the summit of SOX9 bound regions. **B)** Distribution of SOX9 bound regions between TSS/promoter regions (here delimited from 5000 bp upstream to 500 bp downstream of TSS), within gene bodies, or distal (lower panel). Lower panel shows distance from SOX9 bound region and the TSS of the associated genes. **C)** Heatmap of differentially expressed genes with an associated SOX9 bound region. Heatmap shows log2 FPKM values while log2 ratios are displayed in the right panel with selected genes highlighted.

We used GREAT (McLean et al. 2010) to determine genes associated with SOX9 bound regions. Of the associated genes (3094 genes total), 867 were differentially expressed (Fig. 3C), 843 were stably expressed, while 1384 genes were not expressed in either state. Genes with both a SOX9 bound region and differential gene expression were considered direct transcriptional targets of SOX9. Of these, 543 genes (63%) were upregulated by SOX9, while 324 genes (37%) were downregulated. Among the upregulated genes were previously characterized SOX9 target genes, such as ECM genes encoding collagens IX and XI (Oh et al. 2014), as well as other TFs like *SOX5*, *SOX6*, *CREB3L2*, and *NFIA*, highlighting the role of SOX9 as a master regulator (Akiyama et al. 2002; Hino et al. 2014; Garside et al. 2015). Of note, we did not observe SOX9 bound to the enhancer of the well-characterized SOX9 target gene, *COL2A1*, although this gene was highly upregulated with SOX9 expression. Other upregulated target genes include the Homeobox TFs *PRRX1* and *PKNOX2*, both of which have roles in mesenchyme development (Lu et al. 1999; Zhou et al. 2013), while PRRX1 also promotes EMT in cancers (Reichert et al. 2013; Takahashi et al. 2013). Genes possibly repressed by SOX9 include endothelial markers, such as *VWF* and *PECAM1*, as well as genes encoding TFs important for endothelial cell functions, including ERG (Kalna et al. 2019) and ETS2 (Wei et al. 2009).

### SOX9 exhibits different modes of chromatin binding and acts as a pioneer factor at a subset of binding sites in closed chromatin

Since SOX9 primarily bound to distal regulatory elements (enhancers), we questioned which chromatin states allow occupancy by SOX9 and if SOX9 itself is able to alter the chromatin landscape and act as a pioneer TF to reprogram cell fate. To determine the epigenomic impact of SOX9 in HUVECs, we performed ATAC-seq and looked at the CUT&Tag data mentioned above for the enhancer marks H3K4me1, H3K27ac, and H3K27me3 four days after transduction with SOX9 or empty vector. We detected SOX9 bound in both open and closed chromatin regions and in regions with and without pre-existing enhancer marks (Fig. 4A). We classified SOX9 bound regions into four distinct clusters, termed C1-C4, determined by chromatin state with SOX9 bound and in the empty vector control cells. The first cluster, C1 (12%), contains regions with high chromatin accessibility regardless of SOX9 presence, as in the enhancer upstream of the gene encoding the ECM-associated protein CTGF. We also observed high levels of H3K4me1 and H3K27ac in these regions, indicative of active regulatory regions, which increased slightly with SOX9 occupancy. C2 regions (10%) had closed chromatin and no enrichment of the assayed histone marks without SOX9, but with SOX9 bound they gained high chromatin accessibility and enrichment of H3K4me1 and H3K27ac. These newly opened chromatin regions demonstrate pioneer TF activity by SOX9. As shown at the *PRRX1* locus, these regions gained chromatin accessibility and active histone marks with SOX9 binding. Of note, we also detected a C2 type region further upstream of the C1 region of the *CTGF* locus. Both C1 and C2 regions gained increased enrichment of active marks and chromatin accessibility with SOX9 binding, the difference being that C2 regions did not display these active properties at all prior to SOX9 expression and binding (Fig. 4B). C3 (23%) consisted of regions with closed chromatin and no active histone marks but there was high enrichment for the repressive mark H3K27me3, as shown in the *FOXR1* locus. Neither chromatin accessibility, active histone marks, nor H3K27me3 levels were affected by SOX9 binding in these regions. The final C4 cluster, composed of over 50% of SOX9 bound regions, did not show chromatin changes with SOX9 binding and was in quiescent chromatin: closed regions with low levels of histone marks. The locus encoding the ECM enzyme MMP2 is an interesting example as it had a C4 type region in addition to a C2 region and this gene was highly upregulated with SOX9 binding.

**Figure 4.**
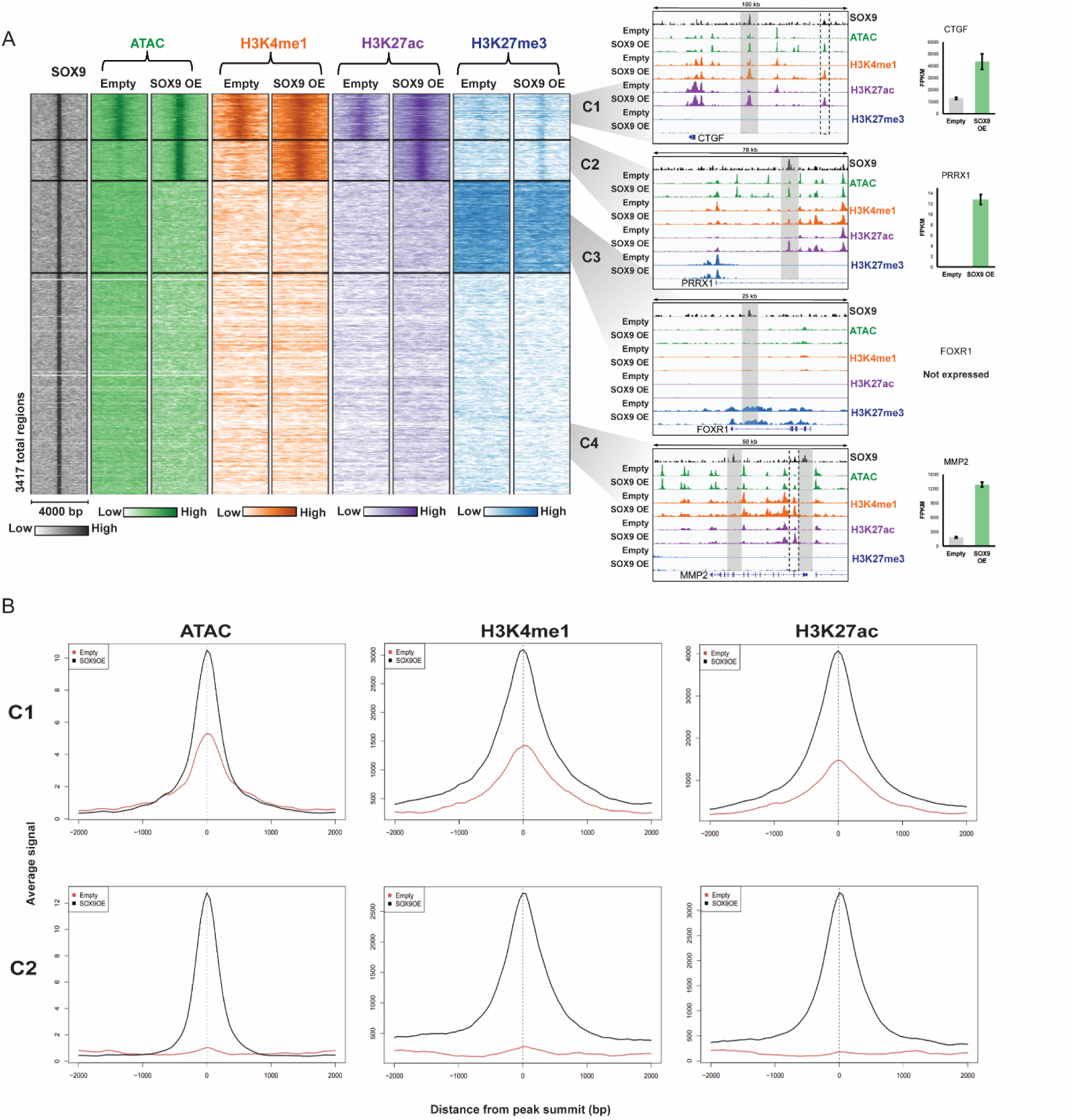
SOX9 opens chromatin and increases enrichment of active histone modifications at a subset of binding sites. **A)** Left panel: heatmap displaying SOX9, ATAC, H3K4me1, H3K27ac, and H3K27me3 signal within a 4 kb window around the summit of SOX9 bound regions. We identified four clusters (C1-C4). The middle panel displays representative loci in each cluster. Regions with a SOX9 peak are highlighted with gray boxes, and the C2 regions in the CTGF and MMP2 loci are highlighted with dashed black boxes. Gene name is upstream of TSS. The right panel shows the expression pattern (average FPKM values) of the given genes. **B)** Average profile plots of ATAC, H3K4me1, and H3K27ac signal within a 4 kb window around the summits of SOX9 bound regions in clusters C1 and C2.

Although most SOX9 binding events occurred in distal regions, we observed C1 had a larger proportion of sites in promoters and/or near the TSS of genes compared to the other clusters (Fig. 5A and S3A). TF binding to chromatin can be categorized by enhanced chromatin occupancy (EChO), a computational strategy recently shown to distinguish direct TF binding to DNA from nucleosomal binding by analyzing average fragment size of called peaks in CUT&RUN data as this reflects sites of minimal DNA protection by proteins (Meers, Janssens, and Henikoff 2019). We used EChO to identify an average fragment size profile in the four clusters. This revealed a smaller average fragment size (<100 bp) for the regions in C1 and C2, reflective of direct DNA binding, and a larger average fragment size (∼150 bp) for C3 and C4 regions, suggesting binding by SOX9 to nucleosomal DNA (Fig. 5B).

**Figure 5.**
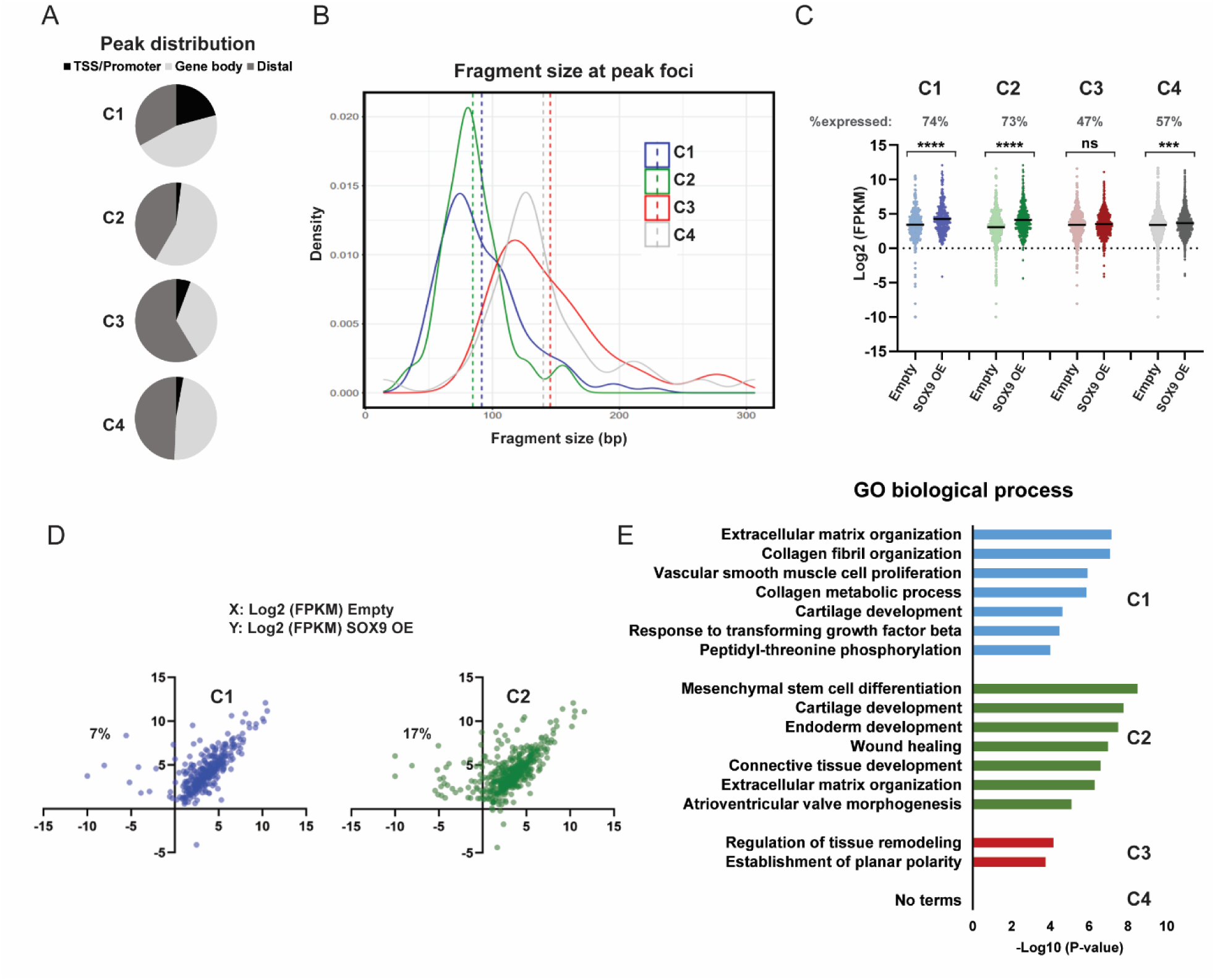
Different modes of SOX9 binding exhibit distinct properties. **A)** Distribution of SOX9 bound regions between TSS/promoter regions (here delimited from 5000 bp upstream to 500 bp downstream of TSS), within gene bodies, or distal (lower panel) in each cluster C1-C4. **B)** Density plot displaying distributions of average fragment size at detected foci for SOX9 CUT&RUN in each cluster C1-C4. Vertical dashed lines represent average fragment size. **C)** Scatterplots of log2 FPKM values of expressed genes (% shown) with an associated SOX9 bound region in each cluster C1-C4. P-values are indicated as in Fig. 1D. Mean is indicated with black line. **D)** Scatterplot comparing the log2 FPKM values between cells transduced with the empty control vector (X-axis) or SOX9 (Y-axis) for all expressed genes in cluster C1 and C2. The percentage of upregulated genes silent in HUVECs transduced with empty vector control that are expressed upon SOX9 expression is indicated. **E)** Top enriched (or the only enriched for C3) biological processes for genes associated with SOX9 bound regions in cluster C1-C4.

Chromatin changes induced by SOX9 binding in C1 and C2 regions lead to an overall upregulation of the associated genes (Fig. 5C, S3B and S3C). There was also upregulation of the genes associated with regions in C4, while expression of genes associated with C3 regions did not significantly change. Of note, many of the upregulated genes associated with C4 regions also had SOX9 binding in C1 or C2 regions (Fig. S3D). After removing all expressed genes that overlap with C1 and/or C2 genes from C4 genes (18%), there was not a significant change in expression of the remaining C4 genes (Fig. S3E and S3F). While over 70% of genes with C1 or C2 type SOX9 binding were expressed, only around 50% of C3 or C4 genes were expressed in either condition (Fig. 5C). As pioneer TFs activate genes in silent chromatin, we asked whether C2 type SOX9 binding, where SOX9 acts as a pioneer TF, is associated with more genes that were silent prior to SOX9 binding compared to the C1 group. This was indeed the case, as 17% of the upregulated genes associated with C2 regions were silenced prior to SOX9 binding, while only 7% of the upregulated C1 genes were silenced before (Fig. 5D).

Genes associated with C1 and C2 regions were enriched for processes connected with mesenchymal cell identity, with C2 regions most enriched for developmental processes and C1 enriched for cellular organization (Fig. 5E). The latter may reflect that C1 genes, such as those encoding ECM components, were already expressed in the endothelial cells prior to SOX9 binding to their active promoters or enhancers, leading to further upregulation. In contrast, genes associated with C2 regions were more specifically linked to a transition to a mesenchymal fate. Notably, we did not identify biological processes associated with the regions in C4, although this was the largest of the four clusters. The latter may indicate that this cluster represents non-specific binding or scanning by SOX9 (Zaret, Lerner, and Iwafuchi-Doi 2016), composing a large group of genes not significantly related to specific biological processes. We observed lower average SOX9 CUT&RUN signal in C3 and C4 regions compared to C1 and C2, suggesting the former are lower-affinity binding events (Fig. S4), although SOX9 must be pausing at these sites to permit detection. Furthermore, we compared SOX9 bound sites with existing whole-genome sequencing datasets, which showed that C1 and C2 regions overlap with binding sites of several TFs, as well as with p300, and RNA polymerase II (Pol2) occupancy, suggesting these are functional regulatory elements (Table S2). Of note, C1 regions overlapped with enhancer elements and active regions (Pol2) in HUVECs, whereas C2 regions mostly overlapped with datasets from cell types where SOX9 is endogenously expressed. In contrast, C3 and C4 regions mostly overlapped with repressed regions in HUVECs and other cell types.

Taken together, our profiling of the chromatin landscape shows that at a subset of enhancers SOX9 binds closed chromatin and increases chromatin accessibility, demonstrating a role as a pioneer TF. Moreover, these enhancers are driving expression of mesenchymal genes and their activation leads to a transition in cell fate.

### Determinants of SOX9 chromatin binding and pioneering activity

As SOX9 binding only resulted in increased chromatin accessibility at a subset of its bound regions in closed chromatin, we questioned if DNA sequence might intrinsically function to dictate how SOX9 binding affects chromatin accessibility and histone marks. Motif analysis revealed that only C2 regions were significantly enriched for SOX motifs (Fig. 6A). Our analysis generated a monomer and a dimer position weight matrix similar to previously identified motifs for SOX9 (Garside et al. 2015; Oh et al. 2014). C3 and C4, where SOX9 binding also occurred in closed chromatin, did not show any evidence of a SOX motif being enriched in the data. This suggested SOX9 binding to closed chromatin, as in C2-C4, was not dictated by a SOX motif, but the ability of SOX9 to open chromatin, as in C2, was dependent on the presence of a SOX motif. However, because the annotated SOX9 monomer and dimer motifs are relatively short and degenerate, respectively, they are found frequently throughout the genome. Therefore, we asked whether there were other unique chromatin features in C2, such as occupancy by other TFs or chromatin modifiers or enrichment of histone variants, which were absent in the other SOX9 bound regions in closed chromatin. When we explored available datasets from HUVECs deposited by the Encyclopedia of DNA Elements (ENCODE) project (Table S3), we discovered that C2 regions were enriched for the histone variant H2A.Z, while C3 and C4 regions were not enriched for any of the chromatin features we examined (Fig. 6B and 6C). Although we observed a lower enrichment of H2A.Z in C2 regions compared to C1 regions (Fig. 6D), the presence of H2A.Z in otherwise silent chromatin may be a determinant for SOX9 pioneering, as we did not observe H2A.Z enriched in C3 and C4 regions. As shown (Fig. 6C), we detected three SOX9 bound regions upstream of the *LGALS3* locus in closed chromatin, but only the region containing H2A.Z became pioneered by SOX9. Furthermore, when we examined all H2A.Z peaks in HUVECs, we observed increased chromatin opening in the regions that also contained a SOX motif with SOX9 expression (Fig. S5A). Although we did not detect SOX9 bound at all sites with H2A.Z enrichment and SOX motifs, these seem to be prerequisites for silent chromatin regions to become opened and activated by SOX9.

**Figure 6.**
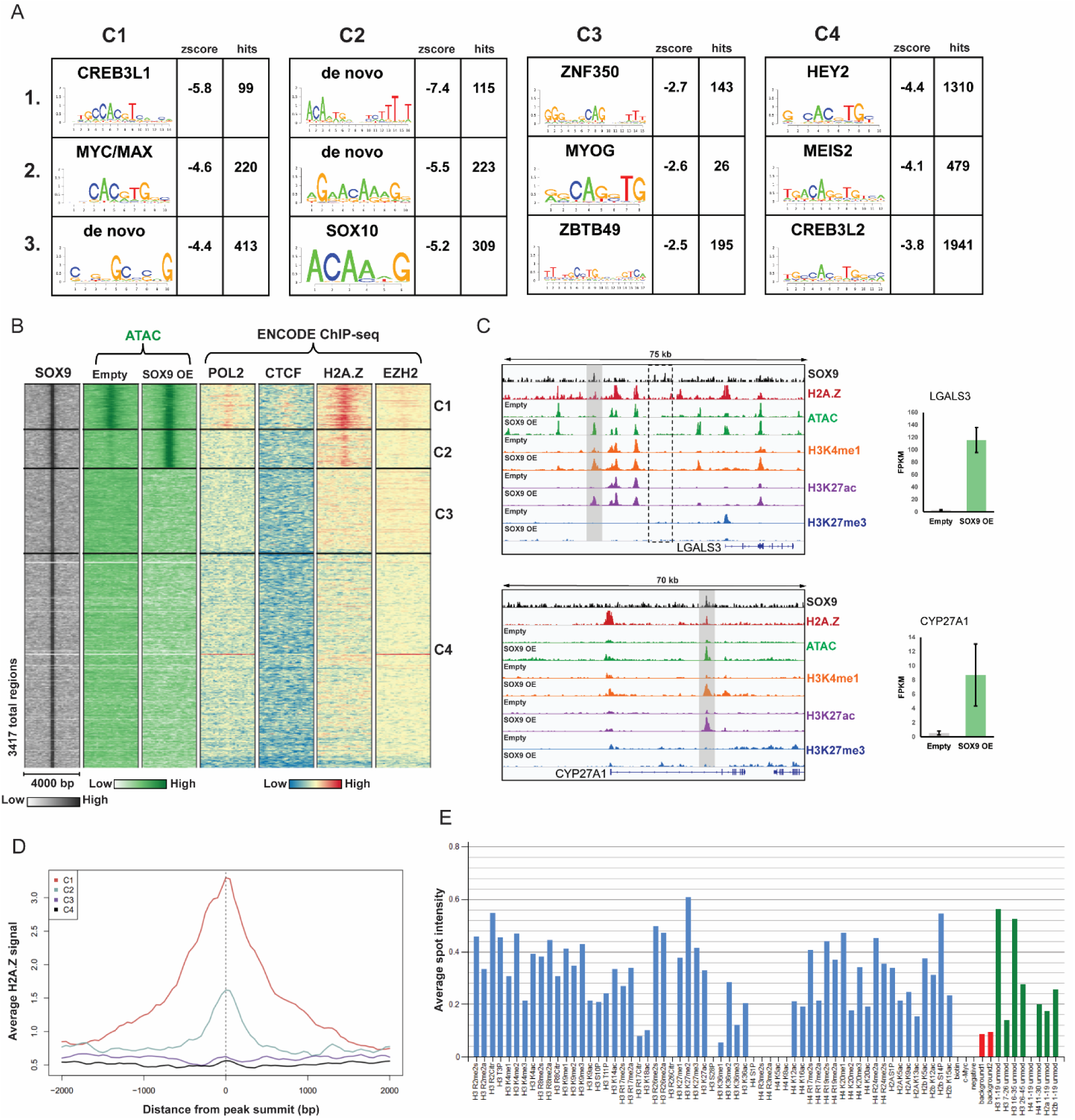
Chromatin features determines SOX9 binding and pioneering activity. **A)** Top three known or de novo TF motifs enriched in SOX9 bound regions in each cluster C1-C4. SOX motifs were only significantly enriched in C2. **B)** Heatmap displaying SOX9, ATAC, RNA Polymerase II (Pol 2) (ENCODE), CTCF (ENCODE), H2A.Z (ENCODE), and EZH2 (ENCODE) signal within a 4 kb window around the summit of SOX9 bound regions. **C)** Representative loci in cluster C2. Regions with SOX9 binding, chromatin opening, and H2A.Z enrichment are highlighted in gray boxes. Regions with SOX9 binding, but no chromatin opening, are highlighted with dashed black boxes. Gene name is upstream of TSS. The right panel shows the expression pattern (average FPKM values) of the given genes. **D)** Average profile plot of H2A.Z signal within a 4 kb window around the summits of SOX9 bound regions in clusters C1-C4. **E)** Relative average spot intensity values for all single modified histone peptides (blue bars), unmodified peptides (green bars), and background array spots (red bars). Histone peptide array was incubated with recombinant SOX9 protein and probed with SOX9 primary antibody.

To further explore chromatin features that may dictate SOX9 binding, we next mapped SOX9 binding to histone tails using a peptide array containing 384 unique histone modification combinations (Fig. S5B and S5C). Strikingly, SOX9 bound most histone tail modifications as well as the unmodified histone tails (Fig. 6E). SOX9 did however not bind to the N-terminal region of H4 and two H3 modifications, H3R26 citrullination and H3S28 phosphorylation, abolished SOX9 histone binding (Fig. 6E and Table S4). We thus concluded that presence of specific histone tail modifications is not a prerequisite for SOX9 to bind chromatin, but certain H3 modifications may impair SOX9 binding and evict SOX9 from chromatin.

### SOX9 binding is dynamic but induces stable changes in the chromatin landscape

To investigate whether there was a global impact on the chromatin landscape regardless of SOX9 binding at a region, we examined all regions that gained or lost chromatin accessibility with SOX9 expression (Fig. 7A and S7A). It is worth noting that most regions that gained accessibility also acquired H3K4me1 and H3K27ac, defining a now active region. Regions with altered chromatin accessibility upon SOX9 expression were exclusively distal and the enriched GO terms for the genes associated with gained regions showed a clear link to EndMT processes, highlighting the role of activated distal enhancers in defining cell-type specificity (Fig. S6A and S6B). Four days after SOX9 transduction, we only detected SOX9 binding at 6% of the total regions with increased chromatin accessibility (C2 regions), suggesting that either other TFs targeted by SOX9 could be responsible for chromatin opening downstream of SOX9 or that SOX9 binding at these regions is transient, allowing SOX9 to bind, chromatin to be opened and activated and SOX9 to move on. Interestingly, after subtraction of all detected SOX9 bound regions, SOX motifs were nonetheless found to be the most significantly enriched in the newly opened chromatin regions (Fig. 7B). SOX dimer motifs were ranked top on our motif list, but a monomer motif was also significantly enriched (Fig. S6C). Regions with decreased chromatin accessibility were enriched for motifs for AP-1 family of TFs, such as JUN and FOS, as well as motifs for the endothelial TFs ETS and ERG (Fig. S7B). Searching for enrichment of overlap between regions with decreased chromatin accessibility and databases of genomic region sets showed overlap with ChIP-seq datasets for AP-1 TFs in HUVECs and DNase hypersensitive regions in endothelial cells (Table S5).

**Figure 7.**
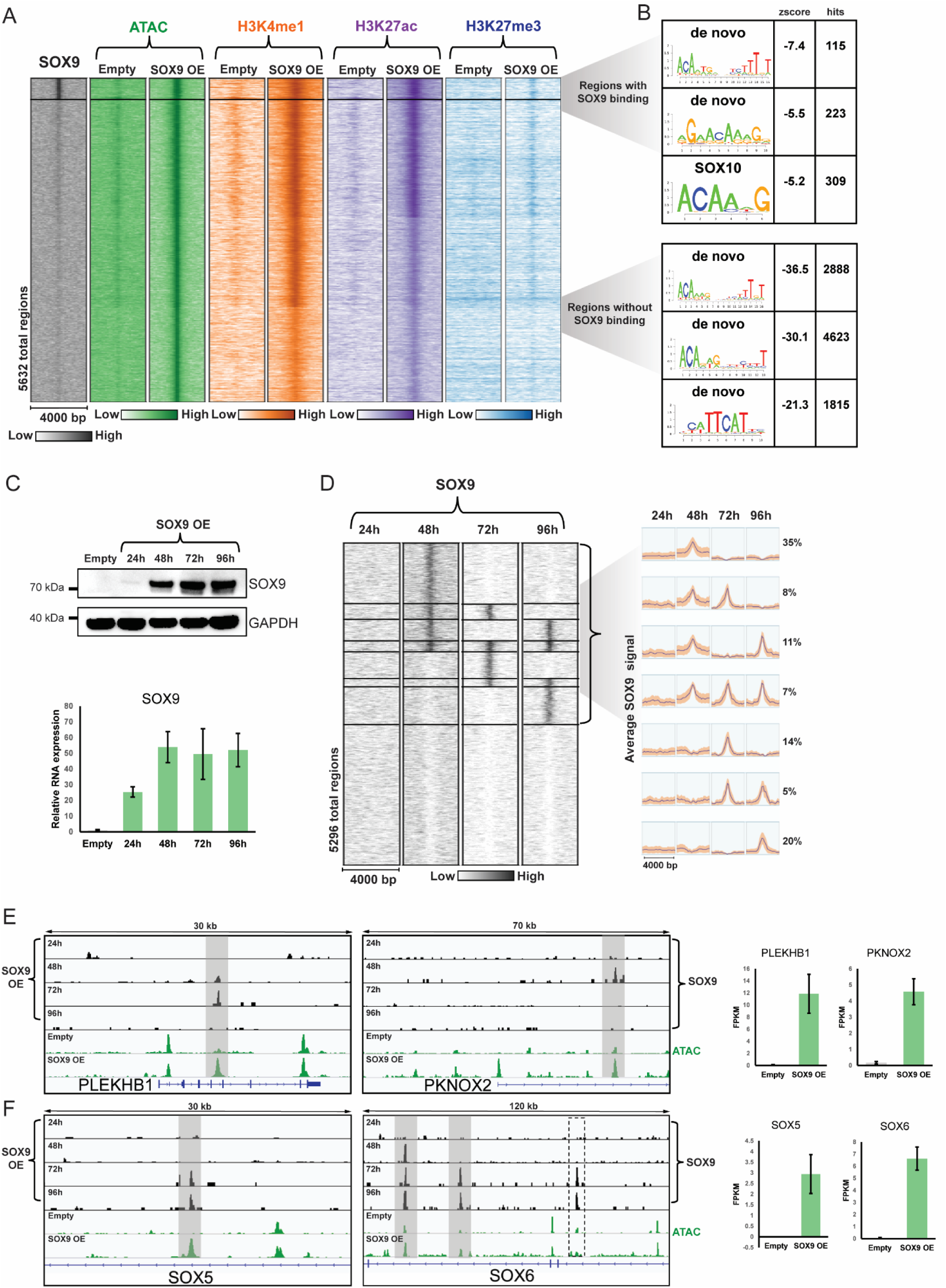
Global changes in chromatin accessibility is caused by dynamic SOX9 binding. **A)** Heatmap displaying SOX9, ATAC, H3K4me1, H3K27ac, and H3K27me3 signal within a 4 kb window around the summit of ATAC peaks in regions with increased chromatin accessibility. The top cluster displays regions with SOX9 binding (C2 from Figure 4A), while the bottom cluster shows regions with increased chromatin accessibility where we did not detect SOX9 binding. **B)** Top three known or de novo TF motifs enriched in regions with increased chromatin accessibility with SOX9 binding (top panel) or without detected SOX9 binding (bottom panel). **C)** Expression of SOX9 (SOX9 OE) confirmed by western blotting and RT-qPCR. Cells were harvested at the indicated timepoints after transduction. **D)** Heatmap displaying SOX9 signal within a 4 kb window around the summit of ATAC peaks in regions with increased chromatin accessibility without detected SOX9 binding in our previous assay. Cells were harvested at the indicated timepoints after transduction. Average SOX9 signal within a 4 kb window around the summit of ATAC peaks in regions with increased chromatin accessibility in regions where we detected SOX9 binding is shown on the right. The percentage of regions with the given SOX9 signal profiles are indicated. **E)** Representative loci where we detected dynamic SOX9 binding and chromatin opening (highlighted in gray boxes). Gene name is upstream of TSS. The right panel shows the expression pattern (average FPKM values) of the given genes. **F)** Representative loci where we detected more stable SOX9 binding and chromatin opening (highlighted in gray boxes). A region with SOX9 binding, but no chromatin opening, is highlighted with a dashed black box. The right panel shows the expression pattern (average FPKM values) of the given genes.

Regions with increased chromatin accessibility showed a similar region overlap as the C2 type SOX9 bound regions, mostly overlapping with ChIP-seq datasets from cells with endogenous SOX9 expression (Table S5). Due to the enrichment of SOX motifs in newly opened chromatin regions where we did not observe SOX9 binding four days post-transduction, we wondered if SOX9 was bound in these regions at earlier timepoints. To determine this, we transduced HUVECs with SOX9 and harvested cells for SOX9 CUT&Tag at four different timepoints post-transduction. SOX9 mRNA and protein was detectable after 24h (Fig. 7C) and by 48h we detected SOX9 CUT&Tag signal in some of the newly opened regions where we did not detect SOX9 bound to in our previous assay (Fig. 7D). Many opened chromatin regions were bound by SOX9 at 48 hours after transduction only, but we observed multiple patterns of SOX9 binding that were strikingly transient and dynamic. In some regions we only observed SOX9 bound at one timepoint (Fig. 7E), while SOX9 bound more stably in other regions (Fig. 7F). Remarkably, SOX9 binding at the tested timepoints accounted for almost 60% of all opening at regions that gained activation. These 2955 regions, in addition to the 336 C2 regions, would all be considered sites where SOX9 acts as a pioneer TF, opening chromatin and encouraging enrichment of active histone marks. Motif analysis of the remaining regions still showed a degenerate SOX dimer motif and a monomer motif among the top enriched motifs, emphasizing transient chromatin binding by SOX9 (Fig. S7D). The chromatin changes induced by SOX9 were, however, stable and driving activation of mesenchymal genes and cell fate transition.

## Discussion

Cell reprogramming in embryonic development is crucial for cell fate and often involves combinations of TFs. Certain TFs, termed pioneer TFs, initiate cell reprogramming events by targeting silent chromatin and TF networks and are thus among the master regulators of cell fate (Iwafuchi-Doi and Zaret 2016). Previous work in mouse has shown that SOX9 is essential in the EndMT process that produces functioning heart valves and acts as a master TF in these cells (Akiyama et al. 2004; Garside et al. 2015). In this study, we used human endothelial cells, HUVECs, to examine whether ectopic SOX9 expression can directly reprogram a mesenchymal cell fate by altering the chromatin landscape. By using genome-wide techniques, we show that SOX9 functions as a pioneer TF by opening chromatin and inducing deposition of active histone marks at previously silent enhancers. The activation of these enhancers drives expression of mesenchymal genes, highlighting the ability of one single TF to induce major chromatin restructuring and cell fate decisions.

Collectively, our analyses provide evidence that ectopic SOX9 expression in HUVECs can drive EndMT by altering the chromatin landscape. SOX9 induced expression of a plethora of TFs associated with mesenchyme, including *PRRX1, PKNOX2*, *FOXC1*, and other SOX TFs, such as *SOX5* and *SOX6*. Interestingly, some TFs with known functions in EndMT/EMT, such as *SNAI1* and *ZEB1*, were downregulated in our dataset. Another factor promoting EMT, *SNAI2*, was not expressed in our dataset. Interestingly, both SNAI1 and SNAI2 are required for EndMT in cardiac cushion morphogenesis in mice (Niessen et al. 2008). Our data may suggest that these factors function upstream of SOX9 or parallel to SOX9 and that SOX9 can initiate EndMT independently of these TFs.

Mapping of SOX9 binding and the chromatin landscape identified different modes of SOX9 binding. Surprisingly, very few SOX9 bound regions localized to open chromatin regions in HUVECs and were rarely found in promoter regions of genes. Previous studies of genome-wide chromatin binding by SOX9 found that 20-30% of SOX9 bound regions occurred within TSS/proximal promoter sites in developing mouse heart valves, limbs, and chondrocytes (Garside et al. 2015; Ohba et al. 2015). In mouse chondrocytes, a SOX motif was enriched in distal regions while promoter regions were enriched for motifs suggesting binding of SOX9 through the basal transcriptional machinery (Ohba et al. 2015). Similarly, in our C1 cluster, where chromatin was open prior to SOX9 binding and where the majority of our promoter bound regions existed, the SOX motif was not significantly enriched, suggesting that at these sites, SOX9 binding may be mediated by other proteins. Furthermore, as previous studies of SOX9 genomic binding assayed endogenous SOX9, SOX9 recruitment to promoters may be dependent on factors absent in HUVECs, possibly explaining why we see less SOX9 bound at promoters.

Pioneer TFs have the unique ability to engage their target sites in closed chromatin, render them accessible and allow other factors to bind (Iwafuchi-Doi 2019). We show that SOX9 harbours this ability but, as observed with other pioneer TFs (Mayran et al. 2018; Takaku et al. 2016; Fuglerud et al. 2017), not all SOX9 binding events result in opening of chromatin. The SOX motif seems to work as an intrinsic signal to guide SOX9 pioneer activity, as pioneering events occurred at regions with SOX dimer and/or monomer motifs. The pluripotency factors OCT4, SOX2, and KLF4,when acting as pioneer TFs, bind nucleosomes *in vitro* by targeting partial DNA motifs displayed on the nucleosome surface (Soufi et al. 2015). According to our data, when SOX9 is acting as a pioneer TF, it requires a full monomer motif or a more degenerate dimer motif (Fig. 6), supporting studies showing that SOX9 chromatin remodeling of nucleosomes assembled *in vitro* was abolished when the SOX motif was disrupted (Coustry et al. 2010).

Pioneering by SOX9 predominantly occurred in regions with enrichment of the histone variant H2A.Z in closed chromatin regions. H2A.Z has been shown to destabilize nucleosomes and is often found at active promoters and enhancers (Martire and Banaszynski 2020), which we also observed in our C1 cluster. The role of H2A.Z in closed chromatin is less clear, but H2A.Z occupancy seems to correlate with pioneer TF binding to nucleosome-occupied regions and has been suggested to act as a scaffold for binding of pioneer TFs and chromatin remodelers (Cauchy, Koch, and Andrau 2017; Subramanian, Fields, and Boyer 2015). Whether SOX9 requires destabilized nucleosomes for chromatin opening and activation of target enhancers remains undetermined, but the requirement for H2A.Z enrichment and motif presence is evident from our study. We could not identify specific histone tail modifications that were required for SOX9 binding. However, modifications in residue 26–28 of canonical H3 that result in less positive charge of the histone tail, namely H3R26cit and H3S28P, strongly impaired SOX9 binding. Of note, methylation of H3R26 did not impact SOX9 binding (compared to the unmodified H3 tail). Both H3R26cit and H3S28P are associated with chromatin decompaction and activation of transcription (Guertin et al. 2014; Sun et al. 2007), suggesting SOX9 is prevented to bind active regulatory regions marked by these modifications.

At sites of SOX9 binding and chromatin opening, we also observed enrichment of the active enhancer marks H3K4me1 and H3K27ac. SOX9 can recruit the histone acetyltransferase p300 to chromatin (Furumatsu et al. 2005), but whether SOX9 also recruits histone methyltransferases to target sites remains unknown. H3K27ac is acquired almost exclusively in the context of pre-existing H3K4me1 at enhancers (Bonn et al. 2012), which suggests that SOX9 may indeed recruit a H3K4 monomethyltransferase to its pioneering target sites. The precise molecular mechanisms underlying pioneer function remain undetermined for most pioneer TFs. Although binding to closed chromatin is a unique property of pioneer TFs, whether pioneer TFs are able to perform chromatin opening fully independently of other factors is still a matter of debate, and the possibility that chromatin remodelers are recruited to sites of pioneer TF binding cannot be ruled out.

Pioneer TF chromatin binding exhibits the potential for low-affinity genome-wide interactions, which may indicate a scanning mechanism (Zaret, Lerner, and Iwafuchi-Doi 2016). SOX9 binding in chromatin that remains repressed and/or quiescent may represent chromatin scanning rather than specific binding by SOX9 (clusters C3 and C4). The lower average SOX9 peak signal at these regions suggests that these are low-affinity binding events and our region set enrichment analysis showed little or no overlap with functional regulatory elements in HUVECs or other cell types (Fig. S4 and Table S2). Since our analysis was performed on a pool of cells, enrichment of SOX9 at the same site suggests there must be other qualities to sites in C3 and C4 that made SOX9 pause at these regions, such as the presence of other chromatin bound factors.

Our time-course SOX9 CUT&Tag revealed that SOX9 chromatin binding is highly dynamic at many of its pioneering sites. However, the chromatin remained open after SOX9 was no longer bound and cell morphology changes persist. The smaller average fragment size of SOX9 bound C2 regions in our EChO analysis indicate that SOX9 does remain bound to DNA after chromatin opening but may be evicted shortly after. Dynamic chromatin binding has been observed for other pioneer TFs (Mayran et al. 2018) and pioneering may be viewed as a one-time event, meaning that SOX9 does not need to remain bound at all sites opened by SOX9 to maintain the chromatin accessibility. Possibly the recruitment of chromatin modifiers and other TFs may act downstream of SOX9 to maintain the open chromatin and ensure stable specification of cell identity.

We observed almost an equal number of enhancers with decreased chromatin accessibility as increased accessibility in HUVECs after SOX9 expression, demonstrating a switch in enhancers initiated by SOX9. The enhancers that were switched off were not bound by SOX9 but were highly enriched for motifs targeted by AP-1 family TFs and ETS family TFs, many of which are highly expressed in HUVEC and downregulated upon SOX9 expression (Fig. S7C). ERG has been found to regulate super-enhancers to promote endothelial homeostasis, while ETS1 has been shown to remodel nucleosomes during T-cell differentiation (Kalna et al. 2019; Cauchy et al. 2016). It is conceivable that ETS family TFs maintain active enhancers in HUVECs, and the downregulation of these factors leads to subsequent loss of chromatin accessibility at their target enhancers.

To summarize, we propose a model whereby SOX9 binds silent chromatin either by transient scanning or by specific binding to regions containing a SOX motif and an H2A.Z-containing nucleosome. In the latter regions, SOX9 opens the local chromatin structure and enriches active histone modifications. In most cases, following chromatin opening, SOX9 gets evicted from the enhancers, but the enhancers remain open and activate associated genes essential for the mesenchymal cell transition. The requirement for SOX9 in regulating the chromatin landscape may differ between cellular systems. For instance, SOX9 may not be crucial for epigenetic changes during chondrogenesis (Liu et al. 2018), but our data shows that expression of SOX9 alone is sufficient to remodel the chromatin landscape for induction of EndMT. In mouse embryonic heart valve development, endocardial cells undergo EndMT at embryonic day 9.5, coinciding with SOX9 expression (Akiyama et al. 2004). The requirement of SOX9 for inducing EndMT in heart valve development has not been clear, but our results indicate that it is sufficient to induce this transition. In adults, tissue damage may stimulate EndMT to give rise to fibroblasts during wound healing or fibrotic diseases. Furthermore, EndMT forms cancer-associated fibroblasts in the tumor microenvironment, which regulates disease progression (Lin, Wang, and Zhang 2012). Thus, SOX9 may be a potential driver of disease mechanisms when aberrantly expressed in endothelial cells.

## Materials and Methods

### Mice

CD1 Elite mice (Charles River) were maintained in accordance with the University of British Columbia’s Animal Care Committee’s standards under specific pathogen-free conditions. Up to five mice were housed per cage and maintained on a 12-hour light-dark cycle. Embryos were collected in ice-cold phosphate-buffered saline (PBS) under a Leica MZ6 dissecting scope. Sex-specific differences were not anticipated and, as such, embryo sex was not determined.

### Immunostaining

For immunostaining, hearts were micro-dissected from E9.5, E10.5, and E12.5 embryos, washed with PBS, and fixed with 4% paraformaldehyde (PFA) at 4°C for 4-12 hours. Ventricles were punctured with fine dissecting forceps prior to fixation at E10.5 and E12.5. After fixation, embryos were washed with PBS three times, then transferred through a sucrose gradient (15%-30%-60% sucrose in PBS, 4°C, 1-12 hours each) before embedding in TissueTek optimal cutting temperature compound. Tissues were frozen on dry-ice, then stored at -80°C until sectioning. 8μm sections were collected using a Leica CM3050S cryostat at -25°C with SuperFrost Plus slides. Sections were circumscribed with an Elite PAP pen (Diagnostic Biosystems K039) and slides were placed in an opaque humidity chamber. Sections were subsequently re-fixed, blocked, stained, and mounted as was done for the HUVEC immunostaining. For HUVEC Immunostaining, cells were seeded on coverslips in 24-well plates six days after transduction. 24 hours later, the cells were washed with PBS and fixed with 4% PFA for 10 minutes at room temperature (RT). After washing in PBS, the slides were incubated in block solution (5% bovine serum albumin, 0.1% Triton-X in PBS) for 1 hour at RT. Slides were then incubated with primary antibodies in block solution overnight at 4 °C. After washing in PBS, the slides were incubated with species-specific Alexa Fluor 568-, 594-, or 647-conjugated secondary antibodies in block solution (all at 1:500) for 1 hour at RT. After washing in PBS, slides were incubated with 1:1000 DAPI to stain nuclei, and then washed again with PBS. Slides were then mounted using a minimal volume of 25mg/mL DABCO in 9:1 glycerol:PBS and fixed in place using clear nail polish. Images were captured using a Nikon Instruments Eclipse Ti confocal laser microscope and image capture and processing was done using Fiji/ImageJ (http://fiji.sc/) with brightness, contrast, and pseudo-coloring adjustments applied equally across all images in a given series. Primary antibodies against the following proteins were used: SOX9 (1:50, AF3045, R&D Systems, and AB5535, Millipore), ERG (1:50, ab92513, Abcam), VIM (1:50, 5741S, Cell Signaling), POSTN (1:50, ab14041, Abcam), PECAM1 (1:50, 102501, Biolegend).

### Cell culture

Primary Human Umbilical Vein Endothelial Cells (HUVECs, Thermo Fisher Scientific) were maintained in Medium 200 supplemented with 2% low serum growth supplement (LSGS, Thermo Fisher Scientific), with penicillin and streptomycin. HUVECs were plated in 6-well plates and lentiviral supernatants were titrated to transduce >95% of the cells (indicated by GFP expression). The media was replaced with fresh Medium 200 24 hours after transduction and cells were collected 72 hours later or seeded for migration assays or immunofluorescence. SOX9 overexpression was confirmed by western blotting and RT-qPCR (described in Supplemental Materials).

### Migration assays

Transwell invasion assays were carried out according to manufacturer’s recommendation. Briefly, 15,000 HUVECs were seeded in the upper chamber of 5 μm 24-well polycarbonate transwell inserts (Corning). HUVECs were maintained for 24 hours under normal culture conditions. After 24 hours, the non-migratory cells were removed from the upper chamber with a cotton swab. The membrane was washed with PBS, fixed with 4% PFA for 10 minutes and stained with DAPI nuclear stain for 10 minutes. The lower side of the membrane was photographed with a Zeiss AxioImager fluorescence microscope. Cells that had migrated through the membrane were counted using Fiji/ImageJ software (http://fiji.sc/). To control for cell death and proliferation, transduced HUVECs of equal density were seeded in 24-wells and monitored for differences after 24h.

### RNA-seq and analysis

RNA was extracted from cells using TRIzol (Thermo Fisher Scientific). Three biological replicates were sequenced by the Biomedical Research Centre (BRC) Sequencing Core at the University of British Columbia. Reads were aligned to Hg19/ GRCh37 using STAR (Dobin et al. 2013). Fragments per kb of exon per million reads (FPKMs) were calculated using Cufflinks (Trapnell et al. 2012). Differential expression was determined by log2 ratio between SOX9 overexpression and empty vector FPKM values. For downstream analyses, only genes with log2 ratio > 0.5 or < -0.5 were included (p-value < 0.05). To determine biological functions and pathways we used Hallmark gene sets (Liberzon et al. 2015).

### CUT&RUN

CUT&RUN was performed as described (Skene, Henikoff, and Henikoff 2018). Briefly, 500,000 cells per experiment were harvested and washed with Wash Buffer (20 mM HEPES pH 7.5, 150 mM NaCl, 0.5 mM spermidine and protease inhibitors), bound to Concanavalin A-coated magnetic beads and incubated with SOX9 antibody (1:100, AB5535, Millipore) diluted in wash buffer containing 0.05% digitonin (Dig-Wash) overnight at 4 °C. Cells were washed and incubated with Guinea Pig anti-Rabbit secondary antibody (1:100, Novus Biologicals) diluted in Dig-Wash for 1 hour at RT. Cells were washed again and incubated with CUTANA™ pAG-MNase (Epicypher) for 10 minutes at RT. Slurry was washed again and placed on ice and incubated with Dig-Wash containing 2 mM CaCl2 for 30 min to activate digestion. Stop buffer (340 mM NaCl, 20 mM EDTA, 4 mM EGTA, 0.05% Digitonin, 0.05 mg/mL glycogen, 5 µg/mL RNase A) was added to stop the reaction, and fragments were released by 30 min incubation at 37 °C. DNA was extracted with phenol-chloroform and ethanol precipitation. Libraries were constructed and sequenced by the BRC Sequencing Core at the University of British Columbia. See Supplemental Material for detailed information about data analysis.

### CUT&Tag

CUT&Tag was performed as described (Kaya-Okur et al. 2019) and pA-Tn5 adapter complex was kindly provided by Steven Henikoff (Fred Hutchinson Cancer Research Center). Cells were harvested, washed, and incubated with primary and secondary antibodies as described for CUT&RUN. Cells were washed again and incubated with pA-Tn5 adapter complex diluted in Dig-300 buffer (0.01% Digitonin, 20 mM HEPES, pH 7.5, 300 mM NaCl, 0.5 mM Spermidine, protease inhibitors) for 1 hour at RT. Cells were washed and incubated with Dig-300 buffer containing 10 mM MgCl2 for 1 hour at 37°C. To stop tagmentation, 16.6 mM EDTA, 0.1% SDS, and 50 μg Proteinase K was added and incubated at 50 °C for 1 hour. DNA was extracted with phenol-chloroform and ethanol precipitation. Libraries were generated using Ad1_noMX and Ad2.3–2.8 barcoded primers from (Buenrostro et al. 2013) and amplified for 14 cycles. DNA purification was carried out using sparQ PureMag Beads (Quantabio). Libraries were sequenced by the BRC Sequencing Core at the University of British Columbia. Primary antibodies against the following proteins were applied: H3K4me1 (1:100, C15410037, Diagenode), H3K27ac (1:100, ab4729, Abcam), H3K27me3 (1:100, C15410069, Diagenode), and SOX9 (1:100, AB5535, Millipore). See Supplemental Material for detailed information about data analysis.

### ATAC-seq

ATAC-seq was performed as described (Buenrostro et al. 2013) with small modifications. 100,000 cells were used per experiment. Libraries were generated using Ad1_noMX and Ad2.1–2.2 barcoded primers from (Buenrostro et al. 2013) and amplified for 10 cycles. DNA was purified using sparQ PureMag Beads (Quantabio) following the manufacturer’s protocol for size selection and library preparation. Libraries were sequenced by the BRC Sequencing Core at the University of British Columbia. See Supplemental Material for detailed information about data analysis.

### Peptide binding assays

MODified™ histone peptide array (13005, Active Motif) was blocked using 3% non-fat milk in TBS. The array was incubated with 2 µg recombinant SOX9 protein (ab131911, Abcam) at 4 °C overnight in CHAPS immunoprecipitation buffer (Fivephoton Biochemicals). After washing in TBST, the array was incubated with SOX9 antibody (1:2000, AB5535, Millipore) in 3% non-fat milk in TBS at 4 °C overnight. After washing in TBST, the array was incubated with HRP-conjugated secondary antibody in 3% non-fat milk in TBS for 1 hour at RT and after washing in TBST, HRP activity was detected with Pierce ECL Western Blotting Substrate (Thermo Fisher Scientific) and imaged using ChemiDoc Imaging System (Bio-Rad). The array was analysed using the manufacturer’s software (Active Motif).

### Data availability

High-throughput sequencing data is publicly available through GEO (accession numbers: GSE155282 and GSE155290).

## Acknowledgements

We are grateful to Steven Henikoff for providing the pA-Tn5 adapter complex for CUT&Tag experiments. B.M.F. is financed by the Research Council of Norway FRIPRO Mobility Research Grant (FRIPRO 275783). P.A.H. receives funding from the Canadian Institutes for Health Research (Funding reference number: 159512).

## Author contributions

B.M.F. and P.A.H. planned the experiments. J.L. dissected and sectioned mouse embryonic hearts and performed immunostaining of hearts and HUVECs. T.L.S. performed transwell migration assays. All other experiments were performed by B.M.F. S.D. performed alignments, processing, and peak-calling on CUT&RUN, CUT&Tag and ATAC-seq data, as well as EChO analysis of the CUT&RUN data. B.M.F. analysed the data. All authors contributed to the writing of the manuscript. B.M.F., J.L., and A.T. made the figures. B.M.F. and P.A.H. obtained funding for the project.

