## Supplemental material for "SOX9 acts as a dynamic pioneer factor inducing stable changes in the chromatin landscape to reprogram endothelial cells"

**Table S1.**
List of genes presented in Figure 2C and supporting references. “Endothelial” genes encode markers of endothelial cells or are genes shown to be downregulated upon EndMT. “Mesenchymal” genes encode mesenchymal markers or have been shown to be upregulated upon EndMT/EMT. Only genes that are expressed in HUVECs with/without SOX9 expression are included.

| Endothelial | Reference |
|---|---|
| ACE | (Nagai et al., 2014) |
| ANTXR1 | (Carson-Walter et al., 2001) |
| CAV1 | (Bauer et al., 2005) |
| CAV2 | (Dragoi et al., 2014) |
| CD151 | (F. Zhang et al., 2011) |
| CD34 | (Pereira et al., 2013) |
| CD93 | (Fonseca et al., 2001) |
| CDH5 | (Pereira et al., 2013) |
| COLEC12 | (Ohtani et al., 2001) |
| CXADR | (Lacher et al., 2006) |
| DCBLD2 | (Nie et al., 2013) |
| ECSCR | (Armstrong et al., 2008) |
| EGFL7 | (Parker et al., 2004) |
| EMCN | (C. Liu et al., 2001) |
| ENG | (Cheifetz et al., 1992) |
| EPOR | (Anagnostou et al., 1994) |
| ERG | (Nikolova-Krstevski et al., 2009) |
| ESAM | (Elcheva et al., 2014) |
| FABP5 | (Masouyé et al., 1997) |
| FLI1 | (Asano et al., 2010) |
| FLT1 | (Seetharam et al., 1995) |
| FLT4 | (Kaipainen et al., 1993) |
| ICAM1 | (Almenar-Queralt et al., 1995) |
| ICAM2 | (Cowan et al., 1998) |
| JUP | (Holen et al., 2012) |
| KDR | (Terman et al., 1991) |
| KLF4 | (Sangwung et al., 2017) |
| KRT19 | (Loriot et al., 2012) |
| LMO2 | (Gratzinger et al., 2009) |
| LYVE1 | (Gordon et al., 2008) |
| MCAM | (Schrage et al., 2008) |
| MTUS1 | (Zhao et al., 2015) |
| NOS3 | (Pereira et al., 2013) |
| PECAM1 | (Pereira et al., 2013) |

| | |
|---|---|
| PODXL | (Horvat et al., 1986) |
| PROCR | (Fukudome & Esmon, 1994) |
| S1PR1 | (Kimura et al., 2000) |
| S1PR3 | (Kimura et al., 2000) |
| SELE | (Collins et al., 1991) |
| SELP | (Polley et al., 1991) |
| SOX18 | (Hosking et al., 2001) |
| SOX7 | (Behrens et al., 2014) |
| STAB1 | (Kzhyshkowska, 2010) |
| TEK | (Pereira et al., 2013) |
| THBD | (Sadler, 1997) |
| THSD1 | (Haasdijk et al., 2016) |
| THSD7A | (C.-H. Wang et al., 2010) |
| TIE1 | (Partanen et al., 1992) |
| TNFRSF10A | (J. H. Li et al., 2003) |
| TNFRSF10B | (Perrot-Applanat et al., 2011) |
| TP53 | (Ghosh et al., 2012) |
| VCAM1 | (Cybulsky & Gimbrone, 1991) |
| VWF | (Pereira et al., 2013) |

| **Mesenchymal** | **Reference** |
|---|---|
| ACTA2 | (Cooley et al., 2014) |
| ADAM12 | (Buenrostro et al., 2015) |
| AHNAK | (Shankar et al., 2010) |
| ALCAM | (Fujiwara et al., 2014) |
| BAMBI | (Fritzmann et al., 2009) |
| BMP2 | (Ma et al., 2005) |
| CALD1 | (Morita et al., 2007) |
| CD44 | (Cho et al., 2012) |
| CDH11 | (Schneider et al., 2012) |
| CDH2 | (M. Wang et al., 2016) |
| CNN1 | (Haque et al., 2011) |
| COL12A1 | (Xiang et al., 2019) |
| COL14A1 | (Aoyagi et al., 2011) |
| COL1A1 | (Gröger et al., 2012) |
| COL1A2 | (Taube et al., 2010) |
| COL3A1 | (Taube et al., 2010) |
| COL5A1 | (Joseph et al., 2014) |
| COL5A2 | (Taube et al., 2010) |

| | |
|---|---|
| COL6A1 | (Jechlinger et al., 2003) |
| COL6A2 | (Jechlinger et al., 2003) |
| COL6A3 | (Huang et al., 2018) |
| COL8A1 | (Minafra et al., 2014) |
| COL9A3 | (Vrljicak et al., 2012) |
| CTGF | (Shafieian et al., 2015) |
| CXCR4 | (X. Li et al., 2014) |
| DDR2 | (Taube et al., 2010) |
| DLC1 | (Taube et al., 2010) |
| FAP | (Taube et al., 2010) |
| FBLN5 | (Taube et al., 2010) |
| FBN1 | (Lien et al., 2019) |
| FOXC1 | (Zhu et al., 2017) |
| HEY1 | (Zavadil et al., 2004) |
| HEY2 | (Lianjie Miao et al., 2018) |
| HEYL | (Bielesz et al., 2010) |
| IGFBP3 | (Gröger et al., 2012) |
| ITGA5 | (Qin et al., 2011) |
| ITGAV | (Wehbe et al., 2012) |
| JAG1 | (Leong et al., 2007) |
| LEF1 | (Kobayashi & Ozawa, 2018) |
| LOXL1 | (Ji et al., 2007) |
| MMP14 | (Sarrió et al., 2008) |
| MMP16 | (Shen et al., 2017) |
| MMP2 | (Wiercinska et al., 2011) |
| MSN | (Haynes et al., 2011) |
| MYH9 | (Beach et al., 2011) |
| NEXN | (Cui et al., 2018) |
| NID2 | (Gröger et al., 2012) |
| NOTCH1 | (Xie et al., 2012) |
| NOTCH3 | (L. Liu et al., 2014) |
| NT5E | (Xiong et al., 2014) |
| P4HA1 | (Gilkes et al., 2013) |
| PLAT | (Yang et al., 2002) |
| PLAU | (Risolino et al., 2014) |
| PLAUR | (Sarrió et al., 2008) |
| POSTN | (Taube et al., 2010) |
| PRKCA | (Gröger et al., 2012) |
| PTX3 | (Taube et al., 2010) |
| RECK | (Gröger et al., 2012) |
| S100A4 | (Mirza et al., 2014) |

| | |
|---|---|
| SERPINE1 | (Risolino et al., 2014) |
| SERPINE2 | (J. Zhang et al., 2020) |
| SLC22A4 | (Gröger et al., 2012) |
| SNAI1 | (Cooley et al., 2014) |
| SPARC | (Fenouille et al., 2012) |
| SPOCK1 | (Liyun Miao et al., 2013) |
| SRF | (Park et al., 2007) |
| SRGN | (Gröger et al., 2012) |
| TAGLN | (Cooley et al., 2014) |
| TGFB1 | (Pardali et al., 2017) |
| TGFB2 | (Medici et al., 2011) |
| TGFBR1 | (Kim et al., 2016) |
| TGFBR2 | (Pino et al., 2010) |
| TPM1 | (Gervasi et al., 2012) |
| TUBA1A | (Gröger et al., 2012) |
| VCAN | (Inai et al., 2013) |
| VIM | (Mendez et al., 2010) |
| WNT5A | (B. Wang et al., 2017) |
| ZEB1 | (Wellner et al., 2009) |
| ZEB2 | (Vandewalle et al., 2005) |

**Table S2.**
The top 20 genomic regions significantly overlapping with SOX9 bound regions in cluster C1-C4 ranked by MeanRnk. The MeanRnk score for each region is the mean rank among three measures: P-value, log odds ratio, and number of overlapping regions. Overlapping regions were identified using LOLA.

| **C1** | | | |
|---|---|---|---|
| **Description** | **Cell type** | **Collection** | **MeanRnk** |
| Enhancers Segments | HUVEC | encode_segmentation | 67 |
| ChIP p300 | SK-N-SH_RA | encode_tfbs | 84.7 |
| ChIP TCF12 | A549 | encode_tfbs | 90 |
| ChIP GR | A549 | encode_tfbs | 92 |
| ChIP c-Myc | MCF10A-Er-Src | encode_tfbs | 98 |
| ChIP c-Fos | HUVEC | encode_tfbs | 103 |
| ChIP Pol2 | SK-N-SH | encode_tfbs | 103 |
| ChIP p300 | HeLa-S3 | encode_tfbs | 113 |
| ChIP FOSL2 | A549 | encode_tfbs | 115 |
| ChIP NR2F2 | Endometrial stromal cell | codex | 115 |
| ChIP GATA-2 | HUVEC | encode_tfbs | 116 |
| ChIP Pol2 | HUVEC | encode_tfbs | 118 |
| ChIP TEAD4 | HepG2 | encode_tfbs | 119 |
| ChIP c-Jun | HUVEC | encode_tfbs | 120 |
| ChIP NR3C1 | A594 cells | cistrome_cistrome | 121 |
| ChIP p300 | A549 | encode_tfbs | 122 |
| ChIP FOXP2 | SK-N-MC | encode_tfbs | 128 |
| ChIP JunD | HeLa-S3 | encode_tfbs | 133 |
| ChIP c-Jun | HeLa-S3 | encode_tfbs | 135 |
| ChIP GR | ECC-1 | encode_tfbs | 137 |

| C2 | | | |
| --- | --- | --- | --- |
| **Description** | **Cell type** | **Collection** | **MeanRnk** |
| DNase HS | Skin cell | sheffield_dnase | 35 |
| ChIP p300 | HepG2 | encode_tfbs | 44.7 |
| ChIP FOXA2 | HepG2 | encode_tfbs | 45 |
| ChIP FOXA1 | HepG2 | encode_tfbs | 51.3 |
| ChIP NFIC | HepG2 | encode_tfbs | 51.3 |
| ChIP SP1 | HepG2 | encode_tfbs | 53.7 |
| ChIP p300 | SK-N-SH_RA | encode_tfbs | 54.3 |
| ChIP HDAC2 | HepG2 | encode_tfbs | 57.3 |
| ChIP TEAD4 | HepG2 | encode_tfbs | 58.7 |
| ChIP RXRA | HepG2 | encode_tfbs | 64 |
| ChIP MYBL2 | HepG2 | encode_tfbs | 65.7 |
| ChIP NR2F2 | Endometrial stromal cell | codex | 66 |
| ChIP ARID3A | HepG2 | encode_tfbs | 67 |
| ChIP HNF4A | HepG2 | encode_tfbs | 75 |
| ChIP TCF7L2 | PANC-1 | encode_tfbs | 77.7 |
| DNase HS | Colo829 | sheffield_dnase | 78 |
| ChIP GR | ECC-1 | encode_tfbs | 78.3 |
| ChIP GR | A549 | encode_tfbs | 78.3 |
| ChIP FOXA1 | A549 | encode_tfbs | 82.7 |
| ChIP NANOG | Embryonic stem cell | codex | 85.3 |

| C3 | | | |
|---|---|---|---|
| **Description** | **Cell type** | **Collection** | **MeanRnk** |
| Repressed Segments | HUVEC | encode_segmentation | 14.7 |
| Repressed Segments | GM12878 | encode_segmentation | 26 |
| ChIP ESR1 | T47D | cistrome_cistrome | 31.3 |
| Repressed Segments | HepG2 | encode_segmentation | 33 |
| Repressed Segments | HeLa-S3 | encode_segmentation | 41 |
| Repressed Segments | K562 | encode_segmentation | 47 |
| ChIP H3K14ac | MCF-7 | cistrome_epigenome | 55.7 |
| ChIP H3K27me3 | MCF-7 | cistrome_epigenome | 56.3 |
| ChIP H3K27me3 | APL164 cells | cistrome_epigenome | 60.7 |
| ChIP CTCF | MCF-7 | encode_tfbs | 61.3 |
| ChIP H3K27me3 | APL164 cells | cistrome_epigenome | 62.7 |
| ChIP H3K9me3 | MCF-7 | cistrome_epigenome | 83.7 |
| ChIP USF-1 | H1-hESC | encode_tfbs | 86.3 |
| DNase HS | Hematopoietic | sheffield_dnase | 89.7 |
| CTCF Segments | H1-hESC | encode_segmentation | 101 |
| ChIP PBX3 | GM12878 | encode_tfbs | 102 |
| ChIP H3K27me3 | NB4 cells | cistrome_epigenome | 107 |
| ChIP ESR1 | MCF-7 | cistrome_cistrome | 107 |
| CTCF Segments | HepG2 | encode_segmentation | 109 |
| UCSC Simple repeats | | ucsc_features | 113 |

| C4 | | | |
|---|---|---|---|
| **Description** | **Cell type** | **Collection** | **MeanRnk** |
| ChIP H3K9me3 | LnCaP | cistrome_epigenome | 43.7 |
| ChIP H3K9me3 | VCaP | cistrome_epigenome | 46.3 |
| ChIP H3K27me3 | UPR9 cells | cistrome_epigenome | 48 |
| UCSC Genomic superdups | | ucsc_features | 50.7 |
| ChIP H3K27me3 | VCaP | cistrome_epigenome | 55.3 |
| ChIP H3K9me3 | Metastatic prostate cancer | cistrome_epigenome | 57.7 |
| ChIP H3K9me3 | NB4 cells | cistrome_epigenome | 63 |
| Transcribed Segments | GM12878 | encode_segmentation | 63.7 |
| ChIP H3K36me3 | LnCaP | cistrome_epigenome | 66 |
| ChIP H3K27me3 | LnCaP | cistrome_epigenome | 68 |
| ChIP Pol2 | Metastatic prostate cancer | cistrome_cistrome | 68.7 |
| Transcribed Segments | HUVEC | encode_segmentation | 69.3 |
| Transcribed Segments | HeLa-S3 | encode_segmentation | 70.7 |
| ChIP H3K9me3 | MCF-7 | cistrome_epigenome | 74 |
| ChIP Pol2 | VCaP | cistrome_cistrome | 77 |
| ChIP H3K9K14ac | APL164 | cistrome_epigenome | 81 |
| ChIP H3K27me3 | Metastatic prostate cancer | cistrome_epigenome | 81 |
| ChIP c-Fos | MCF-7 | cistrome_cistrome | 81.3 |
| ChIP RXR | NB4 cells | cistrome_cistrome | 83.3 |
| ChIP H3K27me3 | MCF-7 | cistrome_epigenome | 86.7 |

**Table S3.**
List of ENCODE datasets (bigwig files) used in Figure 6B.

| HUVEC ChIP dataset | ENCODE Accession |
|---|---|
| POLR2A | ENCFF000YBH |
| CTCF | ENCFF810DGE |
| H2A.Z | ENCFF104GFP |
| EZH2 | ENCFF310GPL |

**Table S4.**

Modifications on histone peptide array incubated with recombinant SOX9 protein and probed with SOX9 antibody ranked by specificity factor. Specificity factor = average intensity of spots on array that contain the modifications/average intensity of spots that do not contain the modification.

| Rank | Modification | Specificity Factor |
|---|---|---|
| 1 | H2A K9ac | 1.6467 |
| 2 | H2A K13ac | 1.6427 |
| 3 | H4 R19me2s | 1.5867 |
| 4 | H2A K5ac | 1.5642 |
| 5 | H2A S1P | 1.4261 |
| 6 | H3 R2me2s | 1.4108 |
| 7 | H3 T3P | 1.4074 |
| 8 | H3 K4me2 | 1.4026 |
| 9 | H3 K9me2 | 1.3528 |
| 10 | H4 R17me2s | 1.351 |
| 11 | H3 R8me2s | 1.3284 |
| 12 | H3 K4ac | 1.3051 |
| 13 | H4 R24me2s | 1.2468 |
| 14 | H3 R2Citr | 1.2423 |
| 15 | H3 K4me1 | 1.2305 |
| 16 | H3 K9me1 | 1.2296 |
| 17 | H3 R2me2a | 1.2278 |
| 18 | H3 K9me3 | 1.22 |
| 19 | H4 R24me2a | 1.1906 |
| 20 | H4 K20me1 | 1.1664 |
| 21 | H3 K4me3 | 1.1574 |
| 22 | H3 K36me2 | 1.1387 |
| 23 | H3 R8Citr | 1.0911 |
| 24 | H3 R8me2a | 1.0442 |
| 25 | H4 R19me2a | 1.0305 |
| 26 | H4 K20me2 | 0.9905 |
| 27 | H3 K27me2 | 0.9752 |
| 28 | H3 T11P | 0.9347 |
| 29 | H4 R17me2a | 0.9344 |
| 30 | H3 K14ac | 0.9067 |
| 31 | H2b K5ac | 0.8957 |
| 32 | H4 K20me3 | 0.8821 |
| 33 | H3 K9ac | 0.8719 |
| 34 | H2b K12ac | 0.8462 |

| | | |
|---|---|---|
| 35 | H3 K36ac | 0.8151 |
| 36 | H3 R17me2a | 0.8034 |
| 37 | H3 R26me2s | 0.7936 |
| 38 | H3 S10P | 0.7886 |
| 39 | H2b S14P | 0.781 |
| 40 | H3 K27me1 | 0.7735 |
| 41 | H4 K20ac | 0.7388 |
| 42 | H3 R26me2a | 0.6951 |
| 43 | H3 K27me3 | 0.617 |
| 44 | H3 R17me2s | 0.5649 |
| 45 | H3 K36me3 | 0.4827 |
| 46 | H2b K15ac | 0.4483 |
| 47 | H4 K16ac | 0.4207 |
| 48 | H3 K27ac | 0.3809 |
| 49 | H3 K18ac | 0.3521 |
| 50 | H3 K36me1 | 0.2108 |
| 51 | H3 R26Citr | 0.1789 |
| 52 | H4 K12ac | 0.1712 |
| 53 | H3 R17Citr | 0.1575 |
| 54 | H4 S1P | 0.011 |
| 55 | H4 R3me2a | 0.011 |
| 56 | H4 R3me2s | 0.006 |
| 57 | H4 K8ac | 0.0059 |
| 58 | H4 K5ac | 0.0032 |
| 59 | H3 S28P | 0 |

**Table S5.**
The top 20 genomic regions significantly overlapping with regions with increased chromatin accessibility or decreased chromatin accessibility upon SOX9 OE. Overlapping regions were identified using LOLA.

| **Increased ATAC** | | | |
|---|---|---|---|
| **Description** | **Cell type** | **Collection** | **MeanRnk** |
| DNase HS | Colo829 | sheffield_dnase | 32.3 |
| DNase HS | Skin cell | sheffield_dnase | 49.3 |
| ChIP TEAD4 | HepG2 | encode_tfbs | 99.3 |
| ChIP p300 | HepG2 | encode_tfbs | 117 |
| ChIP FOXA2 | HepG2 | encode_tfbs | 133 |
| ChIP FOXA1 | HepG2 | encode_tfbs | 135 |
| ChIP NR2F2 | Endometrial stromal cell | codex | 138 |
| ChIP p300 | SK-N-SH_RA | encode_tfbs | 141 |
| ChIP GR | ECC-1 | encode_tfbs | 147 |
| ChIP FOXA1 | A549 | encode_tfbs | 147 |
| ChIP NFIC | HepG2 | encode_tfbs | 151 |
| ChIP SP1 | HepG2 | encode_tfbs | 176 |
| ChIP FOXA1 | ECC-1 | encode_tfbs | 176 |
| ChIP ARID3A | HepG2 | encode_tfbs | 180 |
| ChIP GR | A549 | encode_tfbs | 189 |
| ChIP HDAC2 | HepG2 | encode_tfbs | 190 |
| ChIP ERalpha | ECC-1 | encode_tfbs | 190 |
| ChIP MYBL2 | HepG2 | encode_tfbs | 192 |
| ChIP TCF7L2 | PANC-1 | encode_tfbs | 200 |
| ChIP RXRA | HepG2 | encode_tfbs | 203 |

| Decreased ATAC | | | |
|---|---|---|---|
| **Description** | **Cell type** | **Collection** | **MeanRnk** |
| ChIP c-Fos | HUVEC | encode_tfbs | 4 |
| ChIP c-Jun | HUVEC | encode_tfbs | 18.7 |
| ChIP GATA-2 | HUVEC | encode_tfbs | 31.7 |
| ChIP c-Fos | MCF10A-Er-Src | encode_tfbs | 60 |
| DNase HS | HTR-8 | sheffield_dnase | 68.3 |
| DNase HS | Endothelial | sheffield_dnase | 76.7 |
| ChIP c-Jun | HeLa-S3 | encode_tfbs | 86.3 |
| ChIP STAT3 | MCF10A-Er-Src | encode_tfbs | 86.7 |
| ChIP JunD | HeLa-S3 | encode_tfbs | 91 |
| ChIP c-Fos | Leukaemia cell | codex | 94 |
| ChIP FOSL2 | A549 | encode_tfbs | 99.7 |
| DNase HS | Endothelial;Primitive | sheffield_dnase | 101 |
| ChIP c-FOS | K562 | encode_tfbs | 105 |
| ChIP FOSB | Leukaemia cell | codex | 108 |
| Weak Enhancer | HUVEC | encode_segmentation | 111 |
| ChIP p300 | HeLa-S3 | encode_tfbs | 118 |
| Weak Enhancer | HeLa-S3 | encode_segmentation | 124 |
| ChIP JUNB | K562 | encode_tfbs | 126 |
| ChIP STAT3 | HeLa-S3 | encode_tfbs | 127 |
| ChIP c-Fos | HeLa-S3 | encode_tfbs | 134 |

A

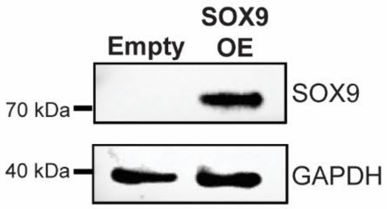


B

| | Empty-1 | Empty-2 | Empty-3 | SOX9-OE-1 | SOX9-OE-2 | SOX9-OE-3 |
|---|---|---|---|---|---|---|
| Empty-1 | 1.000 | 0.990 | 0.989 | 0.893 | 0.879 | 0.893 |
| Empty-2 | 0.990 | 1.000 | 0.989 | 0.882 | 0.874 | 0.885 |
| Empty-3 | 0.989 | 0.989 | 1.000 | 0.896 | 0.886 | 0.904 |
| SOX9-OE-1 | 0.893 | 0.882 | 0.896 | 1.000 | 0.984 | 0.986 |
| SOX9-OE-2 | 0.879 | 0.874 | 0.886 | 0.984 | 1.000 | 0.987 |
| SOX9-OE-3 | 0.893 | 0.885 | 0.904 | 0.986 | 0.987 | 1.000 |

C

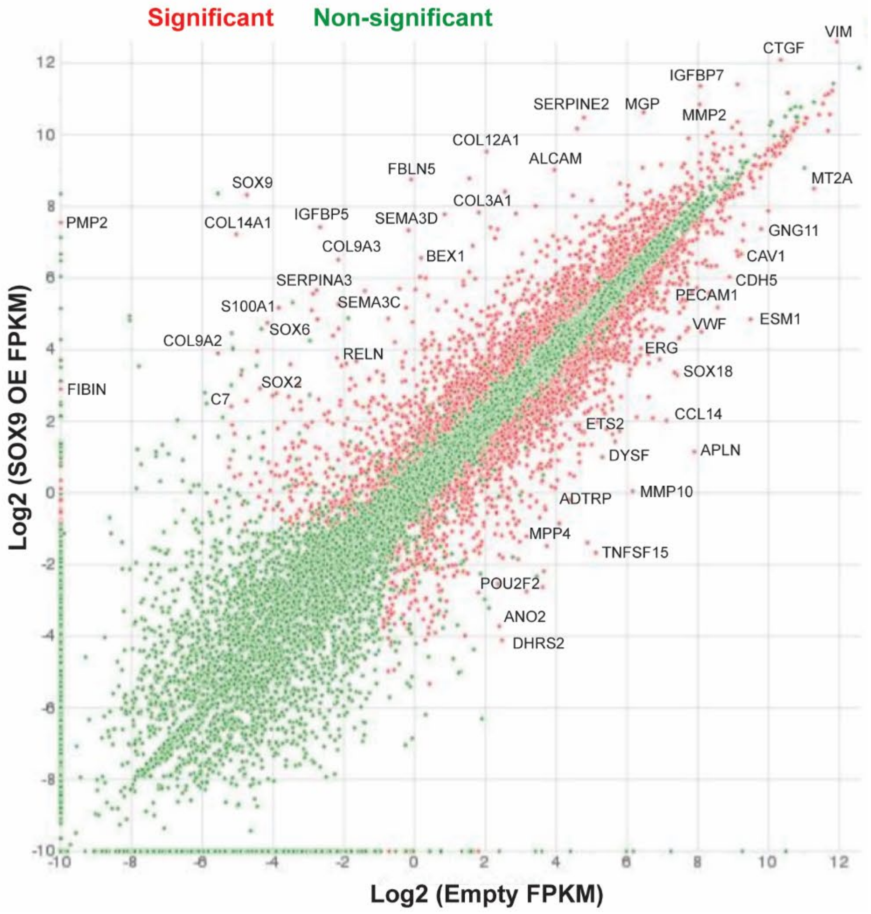

**Figure S1.**

**A)** Expression of SOX9 (SOX9 OE) in transduced HUVECs confirmed by western blotting. **B)** Pearson's correlation plot of FPKM counts visualizing the correlation (r) values between samples. **C)** Scatterplot comparing the normalized average log2 FPKM values between cells transduced with the empty control vector or SOX9 overexpression plasmid. Significantly differentially expressed genes are coloured red with some genes highlighted.

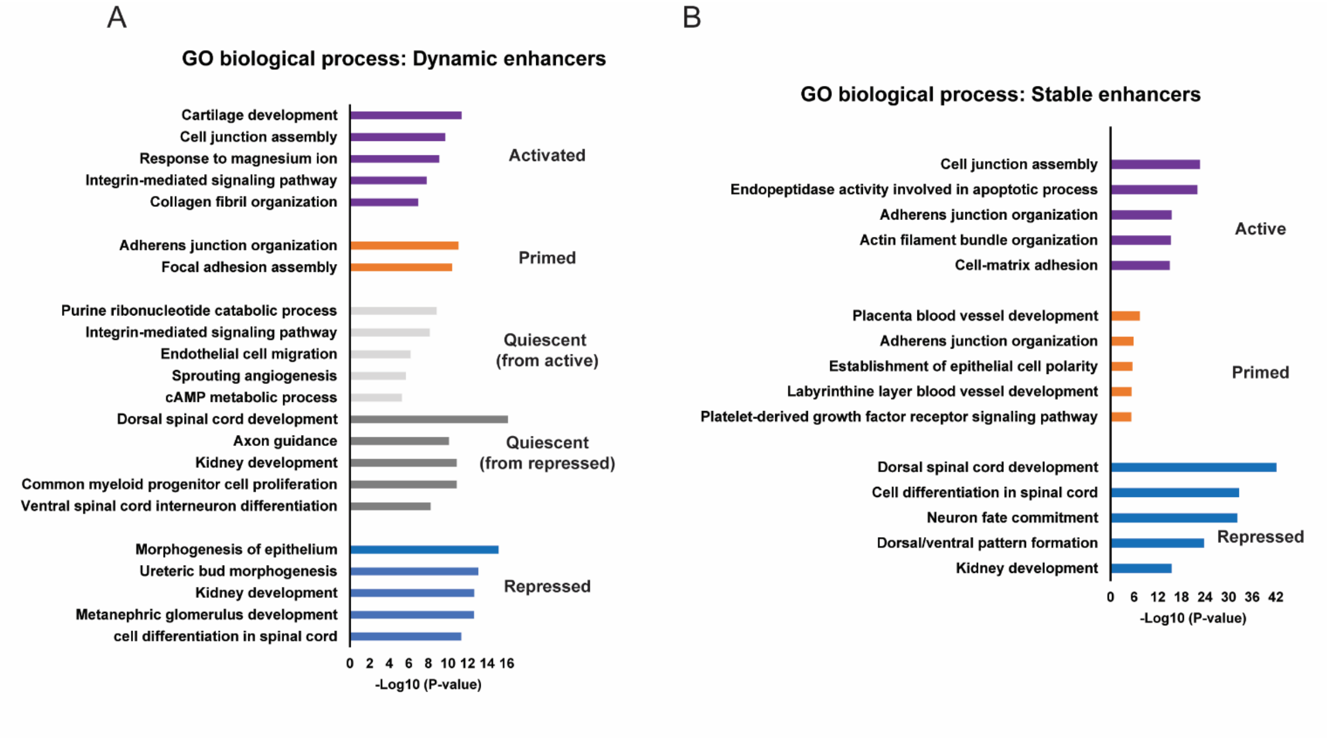


**Figure S2.**
**A)** Top enriched biological processes for genes associated with dynamic enhancers identified in Figure 2. **B)** Top enriched biological processes for genes associated with stable enhancers identified in Figure 2.

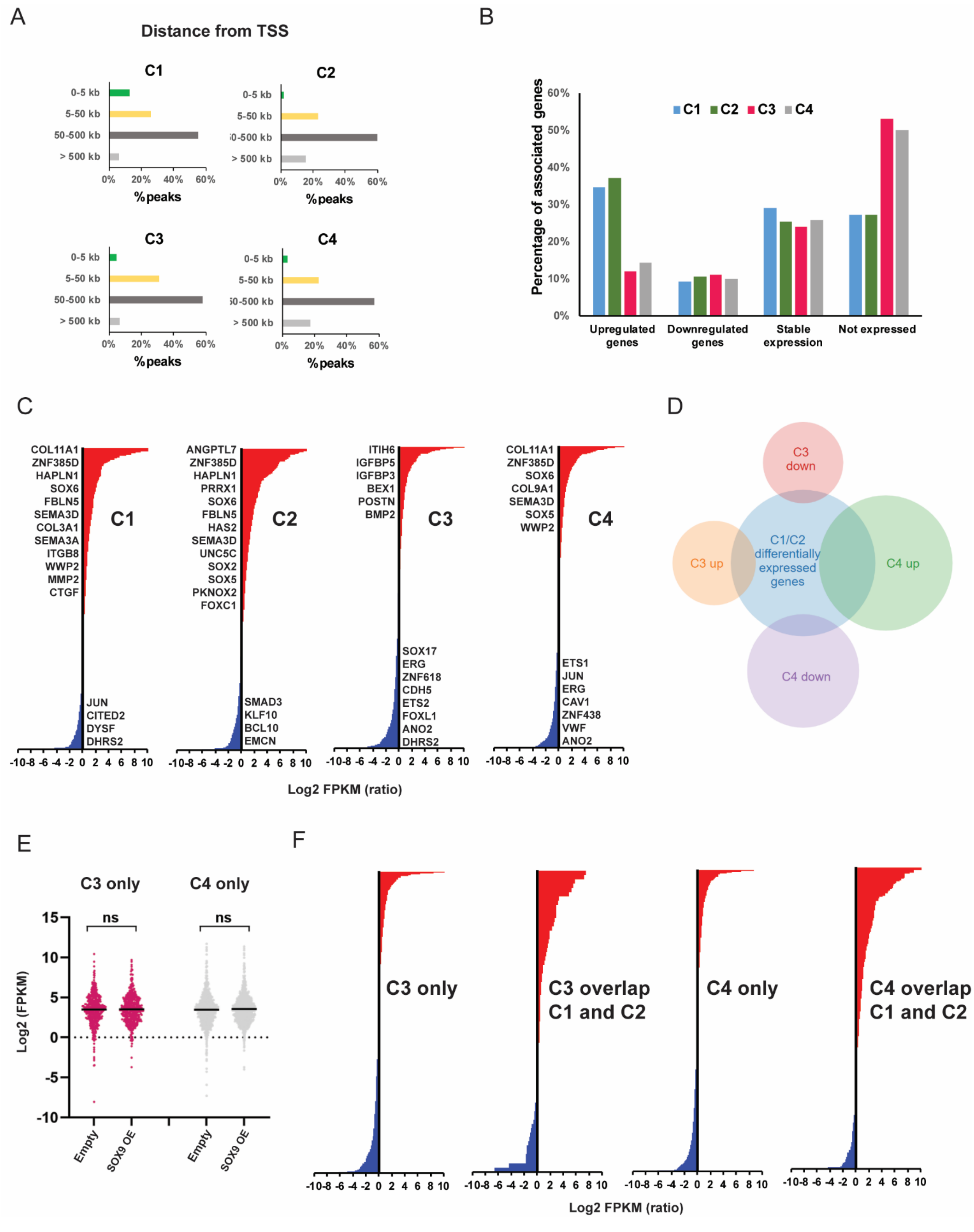


**Figure S3:**

**A)** Distance from SOX9 bound region and the TSS of the associated genes in each cluster C1-C4. **B)**

Percentage of genes associated with SOX9 bound regions in cluster C1-C4 that are upregulated, downregulated, unchanged (stable expression), or not expressed in HUVECs transduced with SOX9. **C)** Log2 FPKM ratios for differentially expressed genes with an associated SOX9 bound region for each cluster C1-C4. Selected genes are highlighted. **D)** Venn diagram showing the number of genes associated with SOX9 bound regions in cluster C3 and C4 that are upregulated or downregulated and their overlap with genes that are differentially expressed and associated with a SOX9 bound region in cluster C1 or C2. **E)** Scatterplots of log2 FPKM values of expressed genes with an associated SOX9 bound region in C3 or C4 which does not overlap with genes in C1 or C2. Significance was evaluated by unpaired, two-tailed *t*-tests with *P*-values (*$P<0.05$; **$P<0.01$; ***$P<0.001$; ****$P<0.0001$; ns $P>0.05$). Mean is indicated with black line. **F)** Log2 FPKM ratios for differentially expressed genes with an associated SOX9 peak found only in C3, C3 overlapping with C1 and C2, only in C4, and C4 overlapping with C1 and C2.

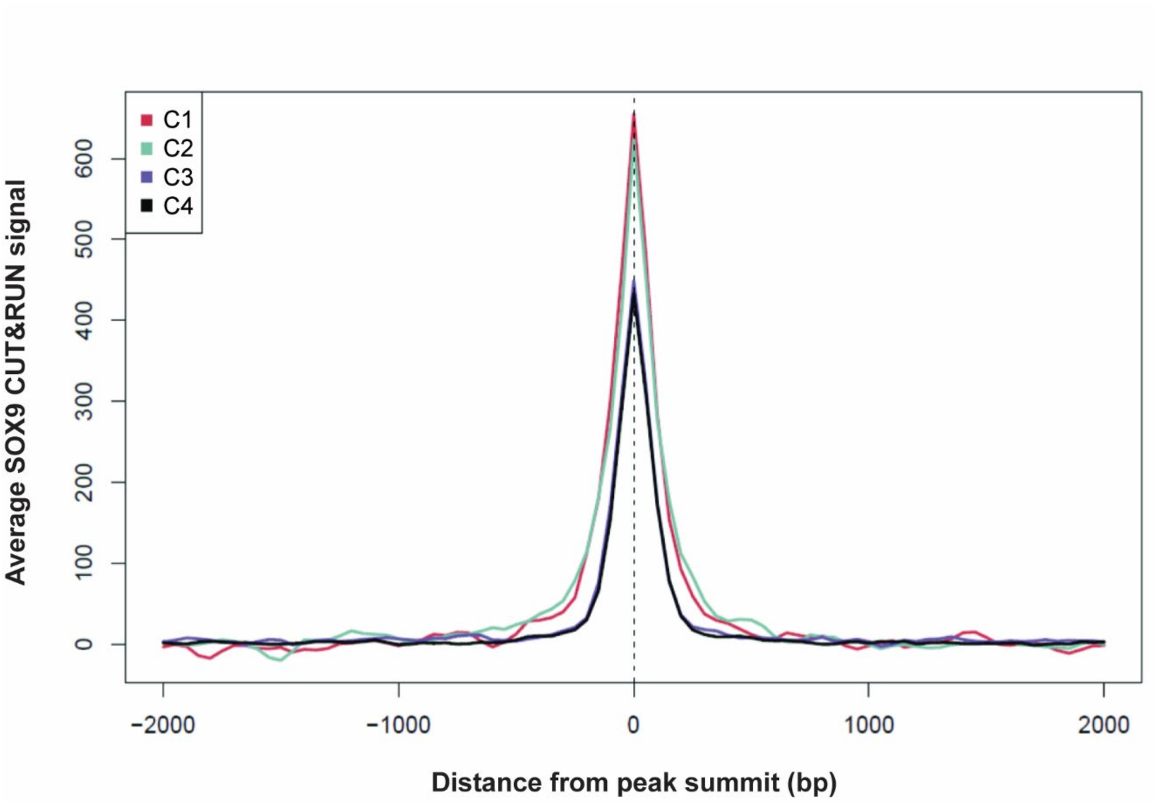


**Figure S4.**
Average profile plot of SOX9 CUT&RUN signal within a 4 kb window around the summits of SOX9 bound regions in each cluster C1-C4.

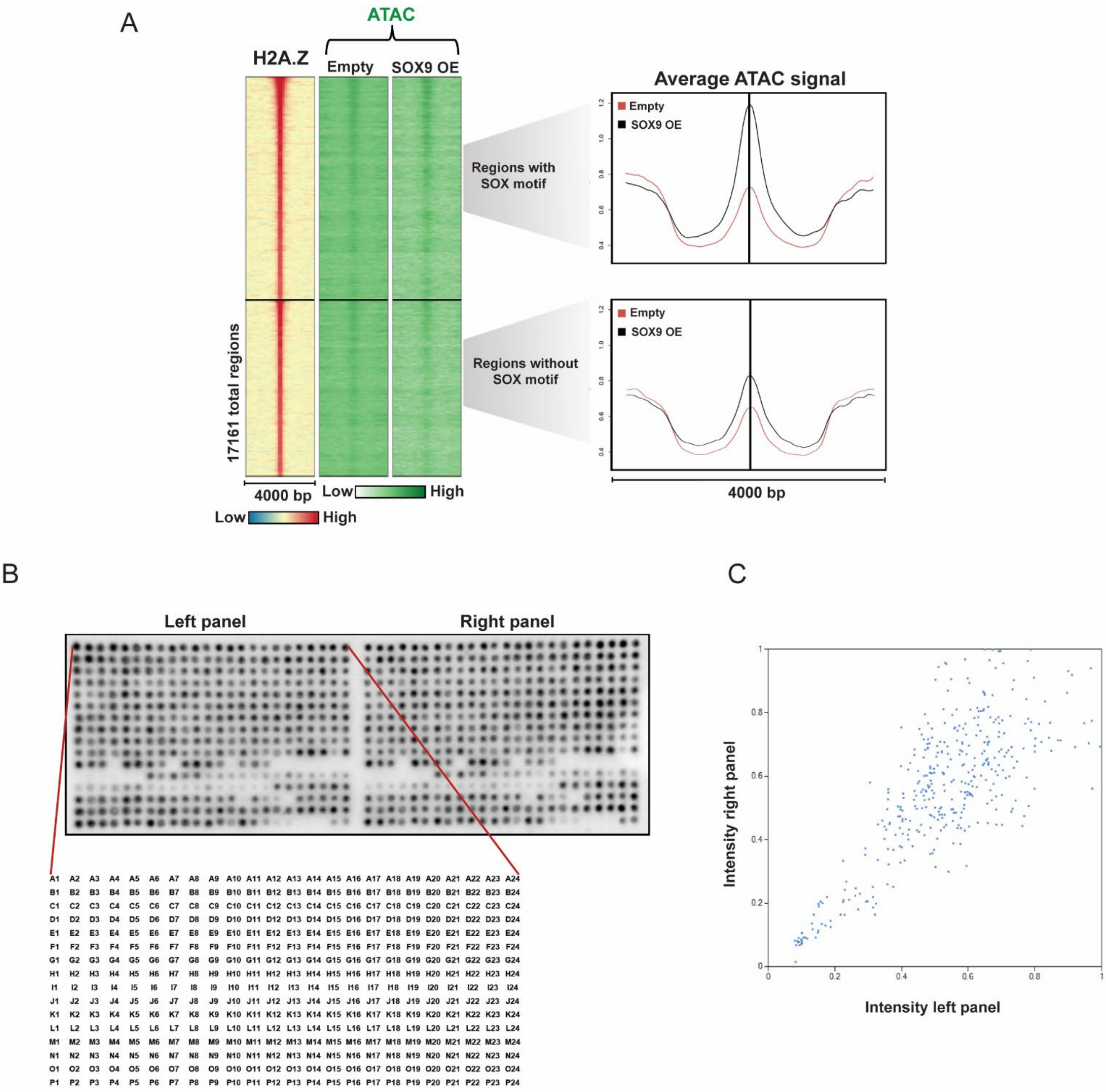


**Figure S5.**
**A)** Left panel:  heatmap displaying H2A.Z ChIP and ATAC signal indicated within a 4 kb window around the summit of H2A.Z peaks. Regions were divided into regions with SOX motifs (monomer and/or dimer) and regions without SOX motifs. The right panel displays the average ATAC signal in H2A.Z regions with SOX motifs or without SOX motifs. **B)** Histone peptide array containing 384 different histone tail modification combinations in duplicate (left panel and right panel) incubated with recombinant SOX9 protein and detected with anti-SOX9 primary antibody. The reference grid for histone peptide locations is shown below. A full overview of the peptide spot positions can be

downloaded from Active Motif's website at www.activemotif.com/modified. **C)** Comparison of the spot intensities on the left and right panel of the histone peptide array in B).

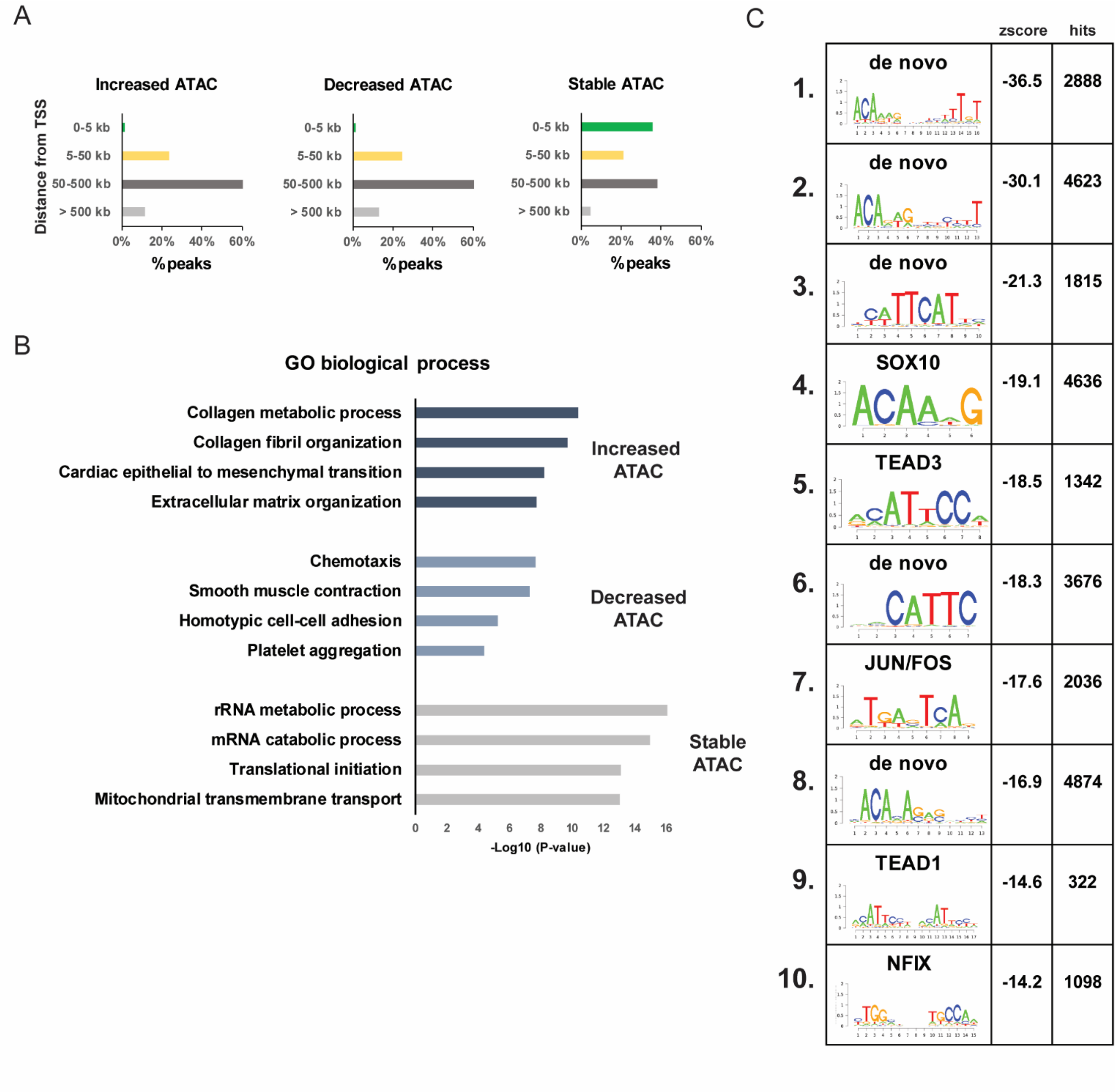


**Figure S6.**
**A)** Distance from regions with increased, decreased, or unchanged (stable) chromatin accessibility (ATAC peaks) and the TSS of the associated genes. **B)** Top enriched biological processes for genes associated with increased, decreased, or unchanged chromatin accessibility regions. **C)** Top ten known or *de novo* TF motifs enriched in regions with increased chromatin accessibility.

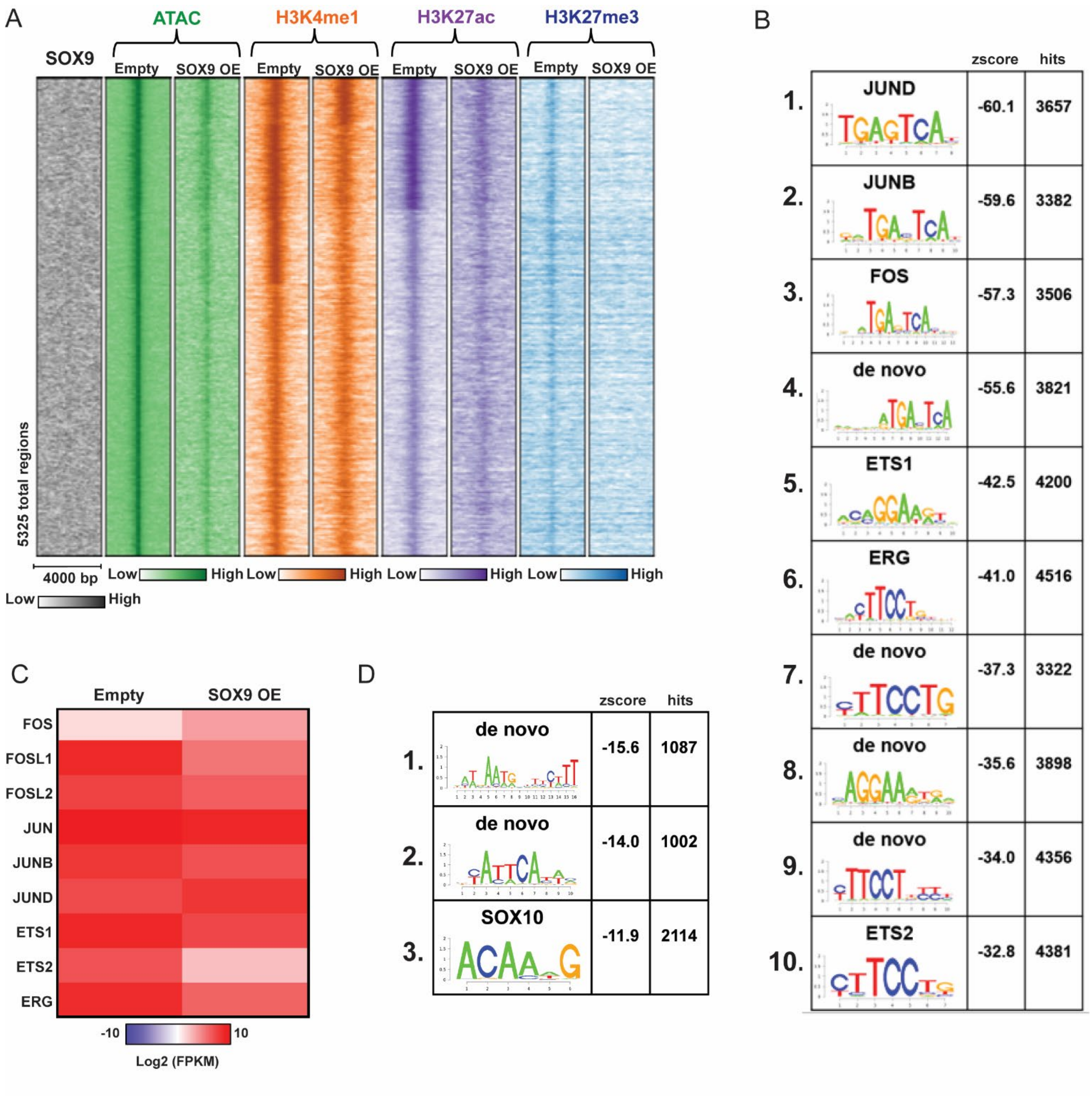


**Figure S7.**

**A)** Heatmap displaying SOX9 CUT&RUN, ATAC, and H3K4me1, H3K27ac, and H3K27me3 CUT&Tag signal indicated within a 4 kb window around the summit of decreased ATAC peaks upon SOX9 expression. **B)** The top ten known or *de novo* TF motifs enriched in regions with decreased chromatin accessibility. **C)** Heatmap displaying log2 FPKM values of expressed TFs with identified motifs in peaks with decreased chromatin accessibility.

### Supplemental Materials and Methods

**Lentiviral vector production.**
Lentivirus empty vector pRRL-cPPT/CTS-MNDU3-PGK-GFP-WPRE and packaging/envelope vectors pCMV-dR8.74, pCMV-VSV-G, and pRSV-Rev were kindly provided by Dr. Andrew Weng (Terry Fox Laboratory, BC Cancer). SOX9 human cDNA was kindly provided by William Stanford (Sprott Centre for Stem Cell Research, Ottawa Hospital Research Institute) and was cloned into pRRL-cPPT/CTS-MNDU3-PGK-GFP-WPRE immediately downstream of the MINDU3 promoter. The construct was verified by sequencing. For lentivirus packaging, HEK293T cells were cultured in Dulbecco's modified Eagle's medium (DMEM) supplemented with 10% fetal bovine serum (FBS). The lentivirus vector and packaging plasmids were cotransfected using polyethyleneimine. Lentiviral supernatants were collected 48 hours after transfection.

**Protein extraction and western blotting.**
SDS loading dye (50 mM Tris-HCl pH 6.8, 2% SDS, 10% glycerol, 12.5 mM EDTA, 0.02% bromophenol blue, 10% beta-mercaptoethanol) was added to pelleted cells and the lysates were sonicated before loaded onto an SDS-polyacrylamide gel and transferred onto a PVDF membrane. The membranes were blocked using 3% non-fat milk in TBS. After blocking, the membranes were incubated overnight at 4 °C with primary antibody. The membranes were then incubated with HRP-conjugated secondary antibody for 1 hour at room temperature. HRP activity was detected with Pierce ECL Western Blotting Substrate (Thermo Fisher Scientific) and imaged using ChemiDoc Imaging System (Bio-Rad). Antibodies against the following proteins were applied: SOX9 (1:2000, AB5535, Millipore) and GAPDH (1:4000, AM4300, Thermo Fisher Scientific).

**RNA isolation, reverse transcription and RT-qPCR.**
RNA was extracted from cells using TRIzol (Thermo Fisher Scientific). 500 ng RNA was used for reverse transcription with First Strand cDNA Synthesis Kit (Roche). RT-qPCR was performed using FastStart Universal SYBR Green Master Mix (Roche) in a StepOnePlus Real-Time PCR System (Applied Biosystems). The values of RNA expression were normalized to the relative amount of the reference gene *GAPDH*. Primer sequences (5'- 3'):

Human GAPDH Forward: GGTGTCGCTGAAGTCAGAG
Human GAPDH Reverse: GGACCTGACCTGCCGTCTAGAA
Human SOX9 Forward: CACGGAGCAGACGCACATCT
Human SOX9 Reverse: TCTCGCTTCAGGTCAGCCTT

**Data processing and analysis of CUT&RUN, CUT&Tag, and ATAC-seq data.**
Reads were mapped to the hg19/GRCh37 genome using the BWA-MEM aligner version 0.7.13 (H. Li and Durbin 2009). The BAM files were filtered using the encode_task_filter.py script from the ENCODE chip-seq-pipeline2 (https://github.com/ENCODE-DCC/chip-seq-pipeline2/blob/master/src/encode_task_filter.py). Briefly, duplicate reads were marked using Picard version 2.1.1 (https://github.com/broadinstitute/picard) and unmapped, unpaired, low quality (MAPQ < 5), non-primary alignment and duplicate reads were removed using SAMtools version 1.3 (H. Li et al. 2009) with the SAM flag filter "-F 1804". MACS2 was used to call peaks using the parameters "-p 0.01

-f BAMPE -g hs" for the CUT&RUN and CUT&Tag data, and "-p 0.01 -g hs --nomodel --shift -75 --extsize 150" for the ATAC-seq data (Zhang et al. 2008). Bigwig files were generated from the filtered BAM files using MACS2 bdgcmp using the fold enrichment option for the ATAC-seq data (Zhang et al. 2008) and deepTools bamCoverage with the options "--binSize 1 --normalizeUsing RPGC --effectiveGenomeSize 2864785220 --extendReads" for the CUT&Tag and CUT&RUN data (Ramírez et al. 2016). All bigwig and bed files were filtered using the ENCODE Blacklist (https://github.com/Boyle-Lab/Blacklist/blob/master/lists/hg19-blacklist.v2.bed.gz). Only peaks with p-value < 0.00001 were considered for further analyses to ensure we only included high-quality peaks. EChO 1.0 was performed as previously described (Meers et al., 2019). Input fragment bed files were generated from prefiltered paired-end BAM files following the steps provided at https://github.com/FredHutch/EChO using samtools 1.6 (H. Li & Durbin, 2009) and bedtools v2.27.1 (Quinlan & Hall, 2010). Bigwig and bed files were analysed using IGV (Robinson et al. 2011), Galaxy (Blankenberg et al. 2010) and Cistrome (T. Liu et al., 2011). Enhancer states were identified using bedtools intersect intervals (Quinlan & Hall, 2010). Differential chromatin accessibility was determined by using bigwigCompare (Ramírez et al. 2016) with 1 kb averaging length. Regions with more than 2-fold increase or decrease in ATAC signal and no called peaks in either the empty vector control cells (for increased chromatin accessibility) or SOX9 OE cells (for decreased chromatin accessibility) were included for further analyses. Motifs were identified using Cistrome's SeqPos (Liu et al. 2011) by scanning 500 bp spanning the SOX9 or ATAC peak summits. K-means clustering of SOX9 binding modes was performed using ChAsE (Younesy et al., 2016). Heatmaps and average profile plots were generated in Cistrome and ChAsE. To identify genes associated with regulatory regions, we used GREAT (McLean et al., 2010). Region set enrichment analysis of SOX9 bound regions and regions with increased or decreased chromatin accessibility was performed using LOLA (Sheffield & Bock, 2016). The regions were tested for enrichment against the LOLA core sequence database.

#### Supplemental References.


Almenar-Queralt, A., Duperray, A., Miles, L. A., Felez, J., & Altieri, D. C. (1995). Apical topography and modulation of ICAM-1 expression on activated endothelium. *The American Journal of Pathology*, *147*(5), 1278–1288.

Anagnostou, A., Liu, Z., Steiner, M., Chin, K., Lee, E. S., Kessimian, N., & Noguchi, C. T. (1994). Erythropoietin receptor mRNA expression in human endothelial cells. *Proceedings of the National Academy of Sciences of the United States of America*, *91*(9), 3974–3978. https://doi.org/10.1073/pnas.91.9.3974

Aoyagi, K., Minashi, K., Igaki, H., Tachimori, Y., Nishimura, T., Hokamura, N., Ashida, A., Daiko, H., Ochiai, A., Muto, M., Ohtsu, A., Yoshida, T., & Sasaki, H. (2011). Artificially induced epithelial-mesenchymal transition in surgical subjects: its implications in clinical and basic cancer research. *PloS One*, *6*(4), e18196. https://doi.org/10.1371/journal.pone.0018196

Armstrong, L.-J., Heath, V. L., Sanderson, S., Kaur, S., Beesley, J. F. J., Herbert, J. M. J., Legg, J. A., Poulsom, R., & Bicknell, R. (2008). ECSM2, an endothelial specific filamin a binding protein that mediates chemotaxis. *Arteriosclerosis, Thrombosis, and Vascular Biology*, *28*(9), 1640–1646. https://doi.org/10.1161/ATVBAHA.108.162511

Asano, Y., Stawski, L., Hant, F., Highland, K., Silver, R., Szalai, G., Watson, D. K., & Trojanowska, M. (2010). Endothelial Fli1 deficiency impairs vascular homeostasis: a role in scleroderma vasculopathy. *The American Journal of Pathology*, *176*(4), 1983–1998. https://doi.org/10.2353/ajpath.2010.090593

Bauer, P. M., Yu, J., Chen, Y., Hickey, R., Bernatchez, P. N., Looft-Wilson, R., Huang, Y., Giordano, F., Stan, R. V, & Sessa, W. C. (2005). Endothelial-specific expression of caveolin-1 impairs microvascular permeability and angiogenesis. *Proceedings of the National Academy of Sciences of the United States of America*, *102*(1), 204–209. https://doi.org/10.1073/pnas.0406092102

Beach, J. R., Hussey, G. S., Miller, T. E., Chaudhury, A., Patel, P., Monslow, J., Zheng, Q., Keri, R. A., Reizes, O., Bresnick, A. R., Howe, P. H., & Egelhoff, T. T. (2011). Myosin II isoform switching mediates invasiveness after TGF-β-induced epithelial-mesenchymal transition. *Proceedings of the National Academy of Sciences of the United States of America*, *108*(44), 17991–17996. https://doi.org/10.1073/pnas.1106499108

Behrens, A. N., Zierold, C., Shi, X., Ren, Y., Koyano-Nakagawa, N., Garry, D. J., & Martin, C. M. (2014). Sox7 is regulated by ETV2 during cardiovascular development. *Stem Cells and Development*, *23*(17), 2004–2013. https://doi.org/10.1089/scd.2013.0525

Bielesz, B., Sirin, Y., Si, H., Niranjan, T., Gruenwald, A., Ahn, S., Kato, H., Pullman, J., Gessler, M., Haase, V. H., & Susztak, K. (2010). Epithelial Notch signaling regulates interstitial fibrosis development in the kidneys of mice and humans. *The Journal of Clinical Investigation*, *120*(11), 4040–4054. https://doi.org/10.1172/JCI43025

Buenrostro, J. D., Wu, B., Chang, H. Y., & Greenleaf, W. J. (2015). ATAC-seq: A Method for Assaying Chromatin Accessibility Genome-Wide. *Current Protocols in Molecular Biology*, *109*, 21.29.1-21.29.9. https://doi.org/10.1002/0471142727.mb2129s109

Carson-Walter, E. B., Watkins, D. N., Nanda, A., Vogelstein, B., Kinzler, K. W., & St Croix, B. (2001). Cell surface tumor endothelial markers are conserved in mice and humans. *Cancer*

*Research*, *61*(18), 6649–6655.

Cheifetz, S., Bellón, T., Calés, C., Vera, S., Bernabeu, C., Massagué, J., & Letarte, M. (1992). Endoglin is a component of the transforming growth factor-beta receptor system in human endothelial cells. *The Journal of Biological Chemistry*, *267*(27), 19027–19030.

Cho, S. H., Park, Y. S., Kim, H. J., Kim, C. H., Lim, S. W., Huh, J. W., Lee, J. H., & Kim, H. R. (2012). CD44 enhances the epithelial-mesenchymal transition in association with colon cancer invasion. *International Journal of Oncology*, *41*(1), 211–218. https://doi.org/10.3892/ijo.2012.1453

Collins, T., Williams, A., Johnston, G. I., Kim, J., Eddy, R., Shows, T., Gimbrone, M. A. J., & Bevilacqua, M. P. (1991). Structure and chromosomal location of the gene for endothelial-leukocyte adhesion molecule 1. *The Journal of Biological Chemistry*, *266*(4), 2466–2473.

Cooley, B. C., Nevado, J., Mellad, J., Yang, D., St Hilaire, C., Negro, A., Fang, F., Chen, G., San, H., Walts, A. D., Schwartzbeck, R. L., Taylor, B., Lanzer, J. D., Wragg, A., Elagha, A., Beltran, L. E., Berry, C., Feil, R., Virmani, R., … Boehm, M. (2014). TGF-β signaling mediates endothelial-to-mesenchymal transition (EndMT) during vein graft remodeling. *Science Translational Medicine*, *6*(227), 227ra34. https://doi.org/10.1126/scitranslmed.3006927

Cowan, P. J., Tsang, D., Pedic, C. M., Abbott, L. R., Shinkel, T. A., d'Apice, A. J., & Pearse, M. J. (1998). The human ICAM-2 promoter is endothelial cell-specific in vitro and in vivo and contains critical Sp1 and GATA binding sites. *The Journal of Biological Chemistry*, *273*(19), 11737–11744. https://doi.org/10.1074/jbc.273.19.11737

Cui, H., Hu, Y., Guo, D., Zhang, A., Gu, Y., Zhang, S., Zhao, C., Gong, P., Shen, X., Li, Y., Wu, H., Wang, L., Zhao, Z., & Fan, H. (2018). DNA methyltransferase 3A isoform b contributes to repressing E-cadherin through cooperation of DNA methylation and H3K27/H3K9 methylation in EMT-related metastasis of gastric cancer. *Oncogene*, *37*(32), 4358–4371. https://doi.org/10.1038/s41388-018-0285-1

Cybulsky, M. I., & Gimbrone, M. A. J. (1991). Endothelial expression of a mononuclear leukocyte adhesion molecule during atherogenesis. *Science (New York, N.Y.)*, *251*(4995), 788–791.

Dragoi, A.-M., Swiss, R., Gao, B., & Agaisse, H. (2014). Novel strategies to enforce an epithelial phenotype in mesenchymal cells. *Cancer Research*, *74*(14), 3659–3672. https://doi.org/10.1158/0008-5472.CAN-13-3231

Elcheva, I., Brok-Volchanskaya, V., Kumar, A., Liu, P., Lee, J.-H., Tong, L., Vodyanik, M., Swanson, S., Stewart, R., Kyba, M., Yakubov, E., Cooke, J., Thomson, J. A., & Slukvin, I. (2014). Direct induction of haematoendothelial programs in human pluripotent stem cells by transcriptional regulators. *Nature Communications*, *5*, 4372. https://doi.org/10.1038/ncomms5372

Fenouille, N., Tichet, M., Dufies, M., Pottier, A., Mogha, A., Soo, J. K., Rocchi, S., Mallavialle, A., Galibert, M.-D., Khammari, A., Lacour, J.-P., Ballotti, R., Deckert, M., & Tartare-Deckert, S. (2012). The epithelial-mesenchymal transition (EMT) regulatory factor SLUG (SNAI2) is a downstream target of SPARC and AKT in promoting melanoma cell invasion. *PloS One*, *7*(7), e40378. https://doi.org/10.1371/journal.pone.0040378

Fonseca, M. I., Carpenter, P. M., Park, M., Palmarini, G., Nelson, E. L., & Tenner, A. J. (2001). C1qR(P), a myeloid cell receptor in blood, is predominantly expressed on endothelial cells in human tissue. *Journal of Leukocyte Biology*, *70*(5), 793–800.

Fritzmann, J., Morkel, M., Besser, D., Budczies, J., Kosel, F., Brembeck, F. H., Stein, U., Fichtner, I., Schlag, P. M., & Birchmeier, W. (2009). A colorectal cancer expression profile that includes transforming growth factor beta inhibitor BAMBI predicts metastatic potential. *Gastroenterology*, *137*(1), 165–175. https://doi.org/10.1053/j.gastro.2009.03.041

Fujiwara, K., Ohuchida, K., Sada, M., Horioka, K., Ulrich, C. D. 3rd, Shindo, K., Ohtsuka, T., Takahata, S., Mizumoto, K., Oda, Y., & Tanaka, M. (2014). CD166/ALCAM expression is characteristic of tumorigenicity and invasive and migratory activities of pancreatic cancer cells. *PloS One*, *9*(9), e107247. https://doi.org/10.1371/journal.pone.0107247

Fukudome, K., & Esmon, C. T. (1994). Identification, cloning, and regulation of a novel endothelial cell protein C/activated protein C receptor. *The Journal of Biological Chemistry*, *269*(42), 26486–26491.

Gervasi, M., Bianchi-Smiraglia, A., Cummings, M., Zheng, Q., Wang, D., Liu, S., & Bakin, A. V. (2012). JunB contributes to Id2 repression and the epithelial-mesenchymal transition in response to transforming growth factor-β. *The Journal of Cell Biology*, *196*(5), 589–603. https://doi.org/10.1083/jcb.201109045

Ghosh, A. K., Nagpal, V., Covington, J. W., Michaels, M. A., & Vaughan, D. E. (2012). Molecular basis of cardiac endothelial-to-mesenchymal transition (EndMT): differential expression of microRNAs during EndMT. *Cellular Signalling*, *24*(5), 1031–1036. https://doi.org/10.1016/j.cellsig.2011.12.024

Gilkes, D. M., Bajpai, S., Chaturvedi, P., Wirtz, D., & Semenza, G. L. (2013). Hypoxia-inducible factor 1 (HIF-1) promotes extracellular matrix remodeling under hypoxic conditions by inducing P4HA1, P4HA2, and PLOD2 expression in fibroblasts. *The Journal of Biological Chemistry*, *288*(15), 10819–10829. https://doi.org/10.1074/jbc.M112.442939

Gordon, E. J., Gale, N. W., & Harvey, N. L. (2008). Expression of the hyaluronan receptor LYVE-1 is not restricted to the lymphatic vasculature; LYVE-1 is also expressed on embryonic blood vessels. *Developmental Dynamics : An Official Publication of the American Association of Anatomists*, *237*(7), 1901–1909. https://doi.org/10.1002/dvdy.21605

Gratzinger, D., Zhao, S., West, R., Rouse, R. V, Vogel, H., Gil, E. C., Levy, R., Lossos, I. S., & Natkunam, Y. (2009). The transcription factor LMO2 is a robust marker of vascular endothelium and vascular neoplasms and selected other entities. *American Journal of Clinical Pathology*, *131*(2), 264–278. https://doi.org/10.1309/AJCP5FP3NAXAXRJE

Gröger, C. J., Grubinger, M., Waldhör, T., Vierlinger, K., & Mikulits, W. (2012). Meta-analysis of gene expression signatures defining the epithelial to mesenchymal transition during cancer progression. *PloS One*, *7*(12), e51136. https://doi.org/10.1371/journal.pone.0051136

Haasdijk, R. A., Den Dekker, W. K., Cheng, C., Tempel, D., Szulcek, R., Bos, F. L., Hermkens, D. M. A., Chrifi, I., Brandt, M. M., Van Dijk, C., Xu, Y. J., Van De Kamp, E. H. M., Blonden, L. A. J., Van Bezu, J., Sluimer, J. C., Biessen, E. A. L., Van Nieuw Amerongen, G. P., & Duckers, H. J. (2016). THSD1 preserves vascular integrity and protects against intraplaque haemorrhaging in ApoE-/- mice. *Cardiovascular Research*, *110*(1), 129–139. https://doi.org/10.1093/cvr/cvw015

Haque, I., Mehta, S., Majumder, M., Dhar, K., De, A., McGregor, D., Van Veldhuizen, P. J., Banerjee, S. K., & Banerjee, S. (2011). Cyr61/CCN1 signaling is critical for epithelial-mesenchymal transition and stemness and promotes pancreatic carcinogenesis. *Molecular Cancer*, *10*, 8.

https://doi.org/10.1186/1476-4598-10-8

Haynes, J., Srivastava, J., Madson, N., Wittmann, T., & Barber, D. L. (2011). Dynamic actin remodeling during epithelial-mesenchymal transition depends on increased moesin expression. *Molecular Biology of the Cell*, *22*(24), 4750–4764. https://doi.org/10.1091/mbc.E11-02-0119

Holen, I., Whitworth, J., Nutter, F., Evans, A., Brown, H. K., Lefley, D. V, Barbaric, I., Jones, M., & Ottewell, P. D. (2012). Loss of plakoglobin promotes decreased cell-cell contact, increased invasion, and breast cancer cell dissemination in vivo. *Breast Cancer Research : BCR*, *14*(3), R86. https://doi.org/10.1186/bcr3201

Horvat, R., Hovorka, A., Dekan, G., Poczewski, H., & Kerjaschki, D. (1986). Endothelial cell membranes contain podocalyxin--the major sialoprotein of visceral glomerular epithelial cells. *The Journal of Cell Biology*, *102*(2), 484–491. https://doi.org/10.1083/jcb.102.2.484

Hosking, B. M., Wang, S. C., Chen, S. L., Penning, S., Koopman, P., & Muscat, G. E. (2001). SOX18 directly interacts with MEF2C in endothelial cells. *Biochemical and Biophysical Research Communications*, *287*(2), 493–500. https://doi.org/10.1006/bbrc.2001.5589

Huang, Y., Li, G., Wang, K., Mu, Z., Xie, Q., Qu, H., Lv, H., & Hu, B. (2018). Collagen Type VI Alpha 3 Chain Promotes Epithelial-Mesenchymal Transition in Bladder Cancer Cells via Transforming Growth Factor β (TGF-β)/Smad Pathway. *Medical Science Monitor : International Medical Journal of Experimental and Clinical Research*, *24*, 5346–5354. https://doi.org/10.12659/MSM.909811

Inai, K., Burnside, J. L., Hoffman, S., Toole, B. P., & Sugi, Y. (2013). BMP-2 induces versican and hyaluronan that contribute to post-EMT AV cushion cell migration. *PloS One*, *8*(10), e77593. https://doi.org/10.1371/journal.pone.0077593

Jechlinger, M., Grunert, S., Tamir, I. H., Janda, E., Lüdemann, S., Waerner, T., Seither, P., Weith, A., Beug, H., & Kraut, N. (2003). Expression profiling of epithelial plasticity in tumor progression. *Oncogene*, *22*(46), 7155–7169. https://doi.org/10.1038/sj.onc.1206887

Ji, H., Ramsey, M. R., Hayes, D. N., Fan, C., McNamara, K., Kozlowski, P., Torrice, C., Wu, M. C., Shimamura, T., Perera, S. A., Liang, M.-C., Cai, D., Naumov, G. N., Bao, L., Contreras, C. M., Li, D., Chen, L., Krishnamurthy, J., Koivunen, J., … Wong, K.-K. (2007). LKB1 modulates lung cancer differentiation and metastasis. *Nature*, *448*(7155), 807–810. https://doi.org/10.1038/nature06030

Joseph, J. V, Conroy, S., Tomar, T., Eggens-Meijer, E., Bhat, K., Copray, S., Walenkamp, A. M. E., Boddeke, E., Balasubramanyian, V., Wagemakers, M., den Dunnen, W. F. A., & Kruyt, F. A. E. (2014). TGF-β is an inducer of ZEB1-dependent mesenchymal transdifferentiation in glioblastoma that is associated with tumor invasion. *Cell Death & Disease*, *5*(10), e1443. https://doi.org/10.1038/cddis.2014.395

Kaipainen, A., Korhonen, J., Pajusola, K., Aprelikova, O., Persico, M. G., Terman, B. I., & Alitalo, K. (1993). The related FLT4, FLT1, and KDR receptor tyrosine kinases show distinct expression patterns in human fetal endothelial cells. *The Journal of Experimental Medicine*, *178*(6), 2077–2088. https://doi.org/10.1084/jem.178.6.2077

Kim, W., Kim, E., Lee, S., Kim, D., Chun, J., Park, K. H., Youn, H., & Youn, B. (2016). TFAP2C-mediated upregulation of TGFBR1 promotes lung tumorigenesis and epithelial-mesenchymal transition. *Experimental & Molecular Medicine*, *48*(11), e273.

https://doi.org/10.1038/emm.2016.125

Kimura, T., Watanabe, T., Sato, K., Kon, J., Tomura, H., Tamama, K., Kuwabara, A., Kanda, T., Kobayashi, I., Ohta, H., Ui, M., & Okajima, F. (2000). Sphingosine 1-phosphate stimulates proliferation and migration of human endothelial cells possibly through the lipid receptors, Edg-1 and Edg-3. *The Biochemical Journal*, *348 Pt 1*(Pt 1), 71–76.

Kobayashi, W., & Ozawa, M. (2018). The epithelial-mesenchymal transition induced by transcription factor LEF-1 is independent of β-catenin. *Biochemistry and Biophysics Reports*, *15*, 13–18. https://doi.org/10.1016/j.bbrep.2018.06.003

Kzhyshkowska, J. (2010). Multifunctional receptor stabilin-1 in homeostasis and disease. *TheScientificWorldJournal*, *10*, 2039–2053. https://doi.org/10.1100/tsw.2010.189

Lacher, M. D., Tiirikainen, M. I., Saunier, E. F., Christian, C., Anders, M., Oft, M., Balmain, A., Akhurst, R. J., & Korn, W. M. (2006). Transforming growth factor-beta receptor inhibition enhances adenoviral infectability of carcinoma cells via up-regulation of Coxsackie and Adenovirus Receptor in conjunction with reversal of epithelial-mesenchymal transition. *Cancer Research*, *66*(3), 1648–1657. https://doi.org/10.1158/0008-5472.CAN-05-2328

Leong, K. G., Niessen, K., Kulic, I., Raouf, A., Eaves, C., Pollet, I., & Karsan, A. (2007). Jagged1-mediated Notch activation induces epithelial-to-mesenchymal transition through Slug-induced repression of E-cadherin. *The Journal of Experimental Medicine*, *204*(12), 2935–2948. https://doi.org/10.1084/jem.20071082

Li, H., & Durbin, R. (2009). Fast and accurate short read alignment with Burrows-Wheeler transform. *Bioinformatics (Oxford, England)*, *25*(14), 1754–1760. https://doi.org/10.1093/bioinformatics/btp324

Li, J. H., Kirkiles-Smith, N. C., McNiff, J. M., & Pober, J. S. (2003). TRAIL induces apoptosis and inflammatory gene expression in human endothelial cells. *Journal of Immunology (Baltimore, Md. : 1950)*, *171*(3), 1526–1533. https://doi.org/10.4049/jimmunol.171.3.1526

Li, X., Li, P., Chang, Y., Xu, Q., Wu, Z., Ma, Q., & Wang, Z. (2014). The SDF-1/CXCR4 axis induces epithelial–mesenchymal transition in hepatocellular carcinoma. *Molecular and Cellular Biochemistry*, *392*(1–2), 77–84. https://doi.org/10.1007/s11010-014-2020-8

Lien, H.-C., Lee, Y.-H., Juang, Y.-L., & Lu, Y.-T. (2019). Fibrillin-1, a novel TGF-beta-induced factor, is preferentially expressed in metaplastic carcinoma with spindle sarcomatous metaplasia. *Pathology*, *51*(4), 375–383. https://doi.org/10.1016/j.pathol.2019.02.001

Liu, C., Shao, Z. M., Zhang, L., Beatty, P., Sartippour, M., Lane, T., Livingston, E., & Nguyen, M. (2001). Human endomucin is an endothelial marker. *Biochemical and Biophysical Research Communications*, *288*(1), 129–136. https://doi.org/10.1006/bbrc.2001.5737

Liu, L., Chen, X., Wang, Y., Qu, Z., Lu, Q., Zhao, J., Yan, X., Zhang, H., & Zhou, Y. (2014). Notch3 is important for TGF-β-induced epithelial-mesenchymal transition in non-small cell lung cancer bone metastasis by regulating ZEB-1. *Cancer Gene Therapy*, *21*(9), 364–372. https://doi.org/10.1038/cgt.2014.39

Liu, T., Ortiz, J. A., Taing, L., Meyer, C. A., Lee, B., Zhang, Y., Shin, H., Wong, S. S., Ma, J., Lei, Y., Pape, U. J., Poidinger, M., Chen, Y., Yeung, K., Brown, M., Turpaz, Y., & Liu, X. S. (2011). Cistrome: an integrative platform for transcriptional regulation studies. *Genome Biology*, *12*(8),

R83. https://doi.org/10.1186/gb-2011-12-8-r83

Loriot, C., Burnichon, N., Gadessaud, N., Vescovo, L., Amar, L., Libé, R., Bertherat, J., Plouin, P.-F., Jeunemaitre, X., Gimenez-Roqueplo, A.-P., & Favier, J. (2012). Epithelial to mesenchymal transition is activated in metastatic pheochromocytomas and paragangliomas caused by SDHB gene mutations. *The Journal of Clinical Endocrinology and Metabolism*, *97*(6), E954-62. https://doi.org/10.1210/jc.2011-3437

Ma, L., Lu, M.-F., Schwartz, R. J., & Martin, J. F. (2005). Bmp2 is essential for cardiac cushion epithelial-mesenchymal transition and myocardial patterning. *Development (Cambridge, England)*, *132*(24), 5601–5611. https://doi.org/10.1242/dev.02156

Masouyé, I., Hagens, G., Van Kuppevelt, T. H., Madsen, P., Saurat, J. H., Veerkamp, J. H., Pepper, M. S., & Siegenthaler, G. (1997). Endothelial cells of the human microvasculature express epidermal fatty acid-binding protein. *Circulation Research*, *81*(3), 297–303. https://doi.org/10.1161/01.res.81.3.297

McLean, C. Y., Bristor, D., Hiller, M., Clarke, S. L., Schaar, B. T., Lowe, C. B., Wenger, A. M., & Bejerano, G. (2010). GREAT improves functional interpretation of cis-regulatory regions. *Nature Biotechnology*, *28*(5), 495–501. https://doi.org/10.1038/nbt.1630

Medici, D., Potenta, S., & Kalluri, R. (2011). Transforming growth factor-β2 promotes Snail-mediated endothelial-mesenchymal transition through convergence of Smad-dependent and Smad-independent signalling. *The Biochemical Journal*, *437*(3), 515–520. https://doi.org/10.1042/BJ20101500

Meers, M. P., Janssens, D. H., & Henikoff, S. (2019). Pioneer Factor-Nucleosome Binding Events during Differentiation Are Motif Encoded. *Molecular Cell*, *75*(3), 562-575.e5. https://doi.org/10.1016/j.molcel.2019.05.025

Mendez, M. G., Kojima, S.-I., & Goldman, R. D. (2010). Vimentin induces changes in cell shape, motility, and adhesion during the epithelial to mesenchymal transition. *FASEB Journal : Official Publication of the Federation of American Societies for Experimental Biology*, *24*(6), 1838–1851. https://doi.org/10.1096/fj.09-151639

Miao, Lianjie, Li, J., Li, J., Tian, X., Lu, Y., Hu, S., Shieh, D., Kanai, R., Zhou, B.-Y., Zhou, B., Liu, J., Firulli, A. B., Martin, J. F., Singer, H., Zhou, B., Xin, H., & Wu, M. (2018). Notch signaling regulates Hey2 expression in a spatiotemporal dependent manner during cardiac morphogenesis and trabecular specification. *Scientific Reports*, *8*(1), 2678. https://doi.org/10.1038/s41598-018-20917-w

Miao, Liyun, Wang, Y., Xia, H., Yao, C., Cai, H., & Song, Y. (2013). SPOCK1 is a novel transforming growth factor-β target gene that regulates lung cancer cell epithelial-mesenchymal transition. *Biochemical and Biophysical Research Communications*, *440*(4), 792–797. https://doi.org/10.1016/j.bbrc.2013.10.024

Minafra, L., Bravatà, V., Forte, G. I., Cammarata, F. P., Gilardi, M. C., & Messa, C. (2014). Gene expression profiling of epithelial-mesenchymal transition in primary breast cancer cell culture. *Anticancer Research*, *34*(5), 2173–2183.

Mirza, A., Foster, L., Valentine, H., Welch, I., West, C. M., & Pritchard, S. (2014). Investigation of the epithelial to mesenchymal transition markers S100A4, vimentin and Snail1 in gastroesophageal junction tumors. *Diseases of the Esophagus : Official Journal of the International Society for*

*Diseases of the Esophagus*, *27*(5), 485–492. https://doi.org/10.1111/j.1442-2050.2012.01435.x

Morita, T., Mayanagi, T., & Sobue, K. (2007). Dual roles of myocardin-related transcription factors in epithelial mesenchymal transition via slug induction and actin remodeling. *The Journal of Cell Biology*, *179*(5), 1027–1042. https://doi.org/10.1083/jcb.200708174

Nagai, T., Kanasaki, M., Srivastava, S. P., Nakamura, Y., Ishigaki, Y., Kitada, M., Shi, S., Kanasaki, K., & Koya, D. (2014). N-acetyl-seryl-aspartyl-lysyl-proline inhibits diabetes-associated kidney fibrosis and endothelial-mesenchymal transition. *BioMed Research International*, *2014*, 696475. https://doi.org/10.1155/2014/696475

Nie, L., Guo, X., Esmailzadeh, L., Zhang, J., Asadi, A., Collinge, M., Li, X., Kim, J.-D., Woolls, M., Jin, S.-W., Dubrac, A., Eichmann, A., Simons, M., Bender, J. R., & Sadeghi, M. M. (2013). Transmembrane protein ESDN promotes endothelial VEGF signaling and regulates angiogenesis. *The Journal of Clinical Investigation*, *123*(12), 5082–5097. https://doi.org/10.1172/JCI67752

Nikolova-Krstevski, V., Yuan, L., Le Bras, A., Vijayaraj, P., Kondo, M., Gebauer, I., Bhasin, M., Carman, C. V, & Oettgen, P. (2009). ERG is required for the differentiation of embryonic stem cells along the endothelial lineage. *BMC Developmental Biology*, *9*, 72. https://doi.org/10.1186/1471-213X-9-72

Ohtani, K., Suzuki, Y., Eda, S., Kawai, T., Kase, T., Keshi, H., Sakai, Y., Fukuoh, A., Sakamoto, T., Itabe, H., Suzutani, T., Ogasawara, M., Yoshida, I., & Wakamiya, N. (2001). The membrane-type collectin CL-P1 is a scavenger receptor on vascular endothelial cells. *The Journal of Biological Chemistry*, *276*(47), 44222–44228. https://doi.org/10.1074/jbc.M103942200

Pardali, E., Sanchez-Duffhues, G., Gomez-Puerto, M. C., & Ten Dijke, P. (2017). TGF-β-Induced Endothelial-Mesenchymal Transition in Fibrotic Diseases. *International Journal of Molecular Sciences*, *18*(10). https://doi.org/10.3390/ijms18102157

Park, M. Y., Kim, K. R., Park, H. S., Park, B.-H., Choi, H. N., Jang, K. Y., Chung, M. J., Kang, M. J., Lee, D. G., & Moon, W. S. (2007). Expression of the serum response factor in hepatocellular carcinoma: implications for epithelial-mesenchymal transition. *International Journal of Oncology*, *31*(6), 1309–1315.

Parker, L. H., Schmidt, M., Jin, S.-W., Gray, A. M., Beis, D., Pham, T., Frantz, G., Palmieri, S., Hillan, K., Stainier, D. Y. R., De Sauvage, F. J., & Ye, W. (2004). The endothelial-cell-derived secreted factor Egfl7 regulates vascular tube formation. *Nature*, *428*(6984), 754–758. https://doi.org/10.1038/nature02416

Partanen, J., Armstrong, E., Mäkelä, T. P., Korhonen, J., Sandberg, M., Renkonen, R., Knuutila, S., Huebner, K., & Alitalo, K. (1992). A novel endothelial cell surface receptor tyrosine kinase with extracellular epidermal growth factor homology domains. *Molecular and Cellular Biology*, *12*(4), 1698–1707. https://doi.org/10.1128/mcb.12.4.1698

Pereira, C.-F., Chang, B., Qiu, J., Niu, X., Papatsenko, D., Hendry, C. E., Clark, N. R., Nomura-Kitabayashi, A., Kovacic, J. C., Ma'ayan, A., Schaniel, C., Lemischka, I. R., & Moore, K. (2013). Induction of a hemogenic program in mouse fibroblasts. *Cell Stem Cell*, *13*(2), 205–218. https://doi.org/10.1016/j.stem.2013.05.024

Perrot-Applanat, M., Vacher, S., Toullec, A., Pelaez, I., Velasco, G., Cormier, F., Saad, H. E. S., Lidereau, R., Baud, V., & Bièche, I. (2011). Similar NF-κB gene signatures in TNF-α treated human endothelial cells and breast tumor biopsies. *PloS One*, *6*(7), e21589.

https://doi.org/10.1371/journal.pone.0021589

Pino, M. S., Kikuchi, H., Zeng, M., Herraiz, M.-T., Sperduti, I., Berger, D., Park, D.-Y., Iafrate, A. J., Zukerberg, L. R., & Chung, D. C. (2010). Epithelial to mesenchymal transition is impaired in colon cancer cells with microsatellite instability. *Gastroenterology*, *138*(4), 1406–1417. https://doi.org/10.1053/j.gastro.2009.12.010

Polley, M. J., Phillips, M. L., Wayner, E., Nudelman, E., Singhal, A. K., Hakomori, S., & Paulson, J. C. (1991). CD62 and endothelial cell-leukocyte adhesion molecule 1 (ELAM-1) recognize the same carbohydrate ligand, sialyl-Lewis x. *Proceedings of the National Academy of Sciences of the United States of America*, *88*(14), 6224–6228. https://doi.org/10.1073/pnas.88.14.6224

Qin, L., Chen, X., Wu, Y., Feng, Z., He, T., Wang, L., Liao, L., & Xu, J. (2011). Steroid receptor coactivator-1 upregulates integrin $\alpha_5$ expression to promote breast cancer cell adhesion and migration. *Cancer Research*, *71*(5), 1742–1751. https://doi.org/10.1158/0008-5472.CAN-10-3453

Quinlan, A. R., & Hall, I. M. (2010). BEDTools: a flexible suite of utilities for comparing genomic features. *Bioinformatics*, *26*(6), 841–842. https://doi.org/10.1093/bioinformatics/btq033

Risolino, M., Mandia, N., Iavarone, F., Dardaei, L., Longobardi, E., Fernandez, S., Talotta, F., Bianchi, F., Pisati, F., Spaggiari, L., Harter, P. N., Mittelbronn, M., Schulte, D., Incoronato, M., Di Fiore, P. P., Blasi, F., & Verde, P. (2014). Transcription factor PREP1 induces EMT and metastasis by controlling the TGF-β-SMAD3 pathway in non-small cell lung adenocarcinoma. *Proceedings of the National Academy of Sciences of the United States of America*, *111*(36), E3775-84. https://doi.org/10.1073/pnas.1407074111

Sadler, J. E. (1997). Thrombomodulin structure and function. *Thrombosis and Haemostasis*, *78*(1), 392–395.

Sangwung, P., Zhou, G., Nayak, L., Chan, E. R., Kumar, S., Kang, D.-W., Zhang, R., Liao, X., Lu, Y., Sugi, K., Fujioka, H., Shi, H., Lapping, S. D., Ghosh, C. C., Higgins, S. J., Parikh, S. M., Jo, H., & Jain, M. K. (2017). KLF2 and KLF4 control endothelial identity and vascular integrity. *JCI Insight*, *2*(4), e91700. https://doi.org/10.1172/jci.insight.91700

Sarrió, D., Rodriguez-Pinilla, S. M., Hardisson, D., Cano, A., Moreno-Bueno, G., & Palacios, J. (2008). Epithelial-mesenchymal transition in breast cancer relates to the basal-like phenotype. *Cancer Research*, *68*(4), 989–997. https://doi.org/10.1158/0008-5472.CAN-07-2017

Schneider, D. J., Wu, M., Le, T. T., Cho, S.-H., Brenner, M. B., Blackburn, M. R., & Agarwal, S. K. (2012). Cadherin-11 contributes to pulmonary fibrosis: potential role in TGF-β production and epithelial to mesenchymal transition. *FASEB Journal : Official Publication of the Federation of American Societies for Experimental Biology*, *26*(2), 503–512. https://doi.org/10.1096/fj.11-186098

Schrage, A., Loddenkemper, C., Erben, U., Lauer, U., Hausdorf, G., Jungblut, P. R., Johnson, J., Knolle, P. A., Zeitz, M., Hamann, A., & Klugewitz, K. (2008). Murine CD146 is widely expressed on endothelial cells and is recognized by the monoclonal antibody ME-9F1. *Histochemistry and Cell Biology*, *129*(4), 441–451. https://doi.org/10.1007/s00418-008-0379-x

Seetharam, L., Gotoh, N., Maru, Y., Neufeld, G., Yamaguchi, S., & Shibuya, M. (1995). A unique signal transduction from FLT tyrosine kinase, a receptor for vascular endothelial growth factor VEGF. *Oncogene*, *10*(1), 135–147.

Shafieian, M., Chen, S., & Wu, S. (2015). Integrin-linked kinase mediates CTGF-induced epithelial to mesenchymal transition in alveolar type II epithelial cells. *Pediatric Research*, *77*(4), 520–527. https://doi.org/10.1038/pr.2015.8

Shankar, J., Messenberg, A., Chan, J., Underhill, T. M., Foster, L. J., & Nabi, I. R. (2010). Pseudopodial actin dynamics control epithelial-mesenchymal transition in metastatic cancer cells. *Cancer Research*, *70*(9), 3780–3790. https://doi.org/10.1158/0008-5472.CAN-09-4439

Sheffield, N. C., & Bock, C. (2016). LOLA: enrichment analysis for genomic region sets and regulatory elements in R and Bioconductor. *Bioinformatics (Oxford, England)*, *32*(4), 587–589. https://doi.org/10.1093/bioinformatics/btv612

Shen, Z., Wang, X., Yu, X., Zhang, Y., & Qin, L. (2017). MMP16 promotes tumor metastasis and indicates poor prognosis in hepatocellular carcinoma. *Oncotarget*, *8*(42), 72197–72204. https://doi.org/10.18632/oncotarget.20060

Taube, J. H., Herschkowitz, J. I., Komurov, K., Zhou, A. Y., Gupta, S., Yang, J., Hartwell, K., Onder, T. T., Gupta, P. B., Evans, K. W., Hollier, B. G., Ram, P. T., Lander, E. S., Rosen, J. M., Weinberg, R. A., & Mani, S. A. (2010). Core epithelial-to-mesenchymal transition interactome gene-expression signature is associated with claudin-low and metaplastic breast cancer subtypes. *Proceedings of the National Academy of Sciences of the United States of America*, *107*(35), 15449–15454. https://doi.org/10.1073/pnas.1004900107

Terman, B. I., Carrion, M. E., Kovacs, E., Rasmussen, B. A., Eddy, R. L., & Shows, T. B. (1991). Identification of a new endothelial cell growth factor receptor tyrosine kinase. *Oncogene*, *6*(9), 1677–1683.

Vandewalle, C., Comijn, J., De Craene, B., Vermassen, P., Bruyneel, E., Andersen, H., Tulchinsky, E., Van Roy, F., & Berx, G. (2005). SIP1/ZEB2 induces EMT by repressing genes of different epithelial cell-cell junctions. *Nucleic Acids Research*, *33*(20), 6566–6578. https://doi.org/10.1093/nar/gki965

Vrljicak, P., Cullum, R., Xu, E., Chang, A. C. Y., Wederell, E. D., Bilenky, M., Jones, S. J. M., Marra, M. A., Karsan, A., & Hoodless, P. A. (2012). Twist1 transcriptional targets in the developing atrio-ventricular canal of the mouse. *PloS One*, *7*(7), e40815. https://doi.org/10.1371/journal.pone.0040815

Wang, B., Tang, Z., Gong, H., Zhu, L., & Liu, X. (2017). Wnt5a promotes epithelial-to-mesenchymal transition and metastasis in non-small-cell lung cancer. *Bioscience Reports*, *37*(6). https://doi.org/10.1042/BSR20171092

Wang, C.-H., Su, P.-T., Du, X.-Y., Kuo, M.-W., Lin, C.-Y., Yang, C.-C., Chan, H.-S., Chang, S.-J., Kuo, C., Seo, K., Leung, L. L., & Chuang, Y.-J. (2010). Thrombospondin type I domain containing 7A (THSD7A) mediates endothelial cell migration and tube formation. *Journal of Cellular Physiology*, *222*(3), 685–694. https://doi.org/10.1002/jcp.21990

Wang, M., Ren, D., Guo, W., Huang, S., Wang, Z., Li, Q., Du, H., Song, L., & Peng, X. (2016). N-cadherin promotes epithelial-mesenchymal transition and cancer stem cell-like traits via ErbB signaling in prostate cancer cells. *International Journal of Oncology*, *48*(2), 595–606. https://doi.org/10.3892/ijo.2015.3270

Wehbe, M., Soudja, S. M., Mas, A., Chasson, L., Guinamard, R., de Tenbossche, C. P., Verdeil, G., Van den Eynde, B., & Schmitt-Verhulst, A.-M. (2012). Epithelial-mesenchymal-transition-like and TGFβ pathways associated with autochthonous inflammatory melanoma development in mice. *PloS One*, *7*(11), e49419. https://doi.org/10.1371/journal.pone.0049419

Wellner, U., Schubert, J., Burk, U. C., Schmalhofer, O., Zhu, F., Sonntag, A., Waldvogel, B., Vannier, C., Darling, D., zur Hausen, A., Brunton, V. G., Morton, J., Sansom, O., Schüler, J., Stemmler, M. P., Herzberger, C., Hopt, U., Keck, T., Brabletz, S., & Brabletz, T. (2009). The EMT-activator ZEB1 promotes tumorigenicity by repressing stemness-inhibiting microRNAs. *Nature Cell Biology*, *11*(12), 1487–1495. https://doi.org/10.1038/ncb1998

Wiercinska, E., Naber, H. P. H., Pardali, E., van der Pluijm, G., van Dam, H., & ten Dijke, P. (2011). The TGF-β/Smad pathway induces breast cancer cell invasion through the up-regulation of matrix metalloproteinase 2 and 9 in a spheroid invasion model system. *Breast Cancer Research and Treatment*, *128*(3), 657–666. https://doi.org/10.1007/s10549-010-1147-x

Xiang, Z., Li, J., Song, S., Wang, J., Cai, W., Hu, W., Ji, J., Zhu, Z., Zang, L., Yan, R., & Yu, Y. (2019). A positive feedback between IDO1 metabolite and COL12A1 via MAPK pathway to promote gastric cancer metastasis. *Journal of Experimental & Clinical Cancer Research : CR*, *38*(1), 314. https://doi.org/10.1186/s13046-019-1318-5

Xie, M., Zhang, L., He, C., Xu, F., Liu, J., Hu, Z., Zhao, L., & Tian, Y. (2012). Activation of Notch-1 enhances epithelial-mesenchymal transition in gefitinib-acquired resistant lung cancer cells. *Journal of Cellular Biochemistry*, *113*(5), 1501–1513. https://doi.org/10.1002/jcb.24019

Xiong, L., Wen, Y., Miao, X., & Yang, Z. (2014). NT5E and FcGBP as key regulators of TGF-1-induced epithelial-mesenchymal transition (EMT) are associated with tumor progression and survival of patients with gallbladder cancer. *Cell and Tissue Research*, *355*(2), 365–374. https://doi.org/10.1007/s00441-013-1752-1

Yang, J., Shultz, R. W., Mars, W. M., Wegner, R. E., Li, Y., Dai, C., Nejak, K., & Liu, Y. (2002). Disruption of tissue-type plasminogen activator gene in mice reduces renal interstitial fibrosis in obstructive nephropathy. *The Journal of Clinical Investigation*, *110*(10), 1525–1538. https://doi.org/10.1172/JCI16219

Younesy, H., Nielsen, C. B., Lorincz, M. C., Jones, S. J. M., Karimi, M. M., & Möller, T. (2016). ChAsE: chromatin analysis and exploration tool. *Bioinformatics (Oxford, England)*, *32*(21), 3324–3326. https://doi.org/10.1093/bioinformatics/btw382

Zavadil, J., Cermak, L., Soto-Nieves, N., & Böttinger, E. P. (2004). Integration of TGF-beta/Smad and Jagged1/Notch signalling in epithelial-to-mesenchymal transition. *The EMBO Journal*, *23*(5), 1155–1165. https://doi.org/10.1038/sj.emboj.7600069

Zhang, F., Michaelson, J. E., Moshiach, S., Sachs, N., Zhao, W., Sun, Y., Sonnenberg, A., Lahti, J. M., Huang, H., & Zhang, X. A. (2011). Tetraspanin CD151 maintains vascular stability by balancing the forces of cell adhesion and cytoskeletal tension. *Blood*, *118*(15), 4274–4284. https://doi.org/10.1182/blood-2011-03-339531

Zhang, J., Luo, A., Huang, F., Gong, T., & Liu, Z. (2020). SERPINE2 promotes esophageal squamous cell carcinoma metastasis by activating BMP4. *Cancer Letters*, *469*, 390–398. https://doi.org/10.1016/j.canlet.2019.11.011

Zhao, T., Ding, X., Chang, B., Zhou, X., & Wang, A. (2015). MTUS1/ATIP3a down-regulation is associated with enhanced migration, invasion and poor prognosis in salivary adenoid cystic carcinoma. *BMC Cancer*, *15*, 203. https://doi.org/10.1186/s12885-015-1209-x

Zhu, X., Wei, L., Bai, Y., Wu, S., & Han, S. (2017). FoxC1 promotes epithelial-mesenchymal transition through PBX1 dependent  transactivation of ZEB2 in esophageal cancer. *American Journal of Cancer Research*, *7*(8), 1642–1653.
